# Distinct transcriptional remodeling of immune and metabolic pathways in severe undernutrition during latent tuberculosis infection does not consistently predict tuberculosis progression across independent cohorts

**DOI:** 10.64898/2026.10.26.756475

**Authors:** Reuben S. Maghembe, Vesla I. Kullaya, Peter M. Mbelele, Ritah Francis Mutagonda, Samweli Bahati, Kennedy M. Ngowi, Lydia Nakiyingi, Megan Null, Jon M. Jacobs, William Evan Johnson, Pranay Sinha, Scott K. Heysell, Stellah George Mpagama, Sana Syed

## Abstract

Undernutrition increases tuberculosis (TB) risk, but whether severe undernutrition during latent TB infection (LTBI) produces blood transcriptional changes that resemble active disease or identify future TB remains unclear. We reanalyzed whole-blood RNA sequencing from adults with LTBI in GSE152218, comparing severe undernutrition (BMI <16 kg/m2; n=16) with adequate nutrition (BMI >=18.5 kg/m2; n=11). Genes and Reactome pathways were evaluated. A seven-module summary was compared with active TB among severely undernourished adults and tested prospectively in GSE79362 and GC6-74/GSE94438 after pathway-representation assessment. No genes reached FDR <0.05 in the primary model; two did only in sensitivity analysis. In contrast, 241 Reactome pathways were significant and directionally concordant in both models (NES rho=0.954), including complement/Fc-receptor, B-cell, phospholipid/phagocytosis, neutrophil, vascular, autophagy, and cytokine-signaling pathways. Most changed opposite to the active-TB comparison (178/241; pathway rho=-0.390). The seven-module score was unrelated to baseline progression in GSE79362 (adjusted OR per SD=1.090; P=0.6922). In GC6, four modules met the prespecified representation criterion; their baseline score was inversely associated with progression (adjusted OR per SD=0.711; P=0.0218), but secondary and longitudinal analyses were non-confirmatory. Severe undernutrition during LTBI is associated with reproducible immune- and metabolism-related transcriptional remodeling that differs from active-TB biology and does not consistently identify future TB progression.

## 1. Introduction

Tuberculosis (TB) remains a leading infectious cause of mortality. Undernutrition is a major modifiable driver of incident TB and is associated with poorer outcomes [1-4]. The relationship is bidirectional: undernutrition can increase TB vulnerability, whereas active disease can worsen nutrition through reduced intake and inflammation-associated catabolism [2,4].

Whole-blood RNA sequencing can reveal coordinated transcriptional changes in immune and metabolic processes but does not directly measure proteins, cytokines, metabolites, or metabolic flux. Here, immune and metabolic pathways therefore refer to RNA-derived pathway enrichment.

Blood transcriptomic studies have identified interferon-inducible and neutrophil-linked signatures in active TB and signatures associated with near-term incident disease [7-10], but whether severe undernutrition during LTBI produces the same pattern or consistently predicts progression is unresolved.

Studies directly examining undernutrition and TB immune biology remain limited. Johnson et al. [11] evaluated TB gene-signature behavior in malnourished individuals, and VanValkenburg et al. [12] reported increased inflammatory and TB-risk-signature activity in malnourished household contacts with LTBI. More recently, Locks et al. [32] reported attenuated M. tuberculosis-specific Th1 responses in undernourished household contacts with partial reconstitution after nutritional support. These observations establish a strong rationale for nutrition-aware transcriptomics but do not show whether a severe-undernutrition-associated transcriptional state resembles active-TB biology or behaves consistently as a prospective progression signal.

We therefore asked three linked questions using public whole-blood transcriptomic data. First, what gene- and pathway-level transcriptional changes distinguish severe from adequate nutritional status among adults with LTBI in GSE152218? Second, do those changes resemble the active-TB-versus-LTBI difference among severely undernourished adults in the same dataset? Third, does a seven-module summary of the undernutrition-associated pattern identify future TB in two independent prospective cohorts, GSE79362 and Grand Challenges 6-74 (GC6-74; GSE94438), after accounting for differences in measured gene coverage?

## 2. Methods

### 2.1. Study design and source cohorts

This secondary analysis used three publicly available whole-blood transcriptomic datasets with distinct roles. GSE152218 was used for severe-undernutrition discovery among adults with LTBI and for an exploratory active-TB-versus-LTBI comparison among severely undernourished adults. GSE79362 and GSE94438/GC6 were independent prospective cohorts used to test the discovery-derived seven-module score [9,10,12-15]. The seven modules were selected before external progression outcomes were examined. Extended procedures are provided in the Supplementary Methods (Supplementary material 1).

Figure 1 summarizes the study design and the distinct role of each cohort.

**Figure 1.**
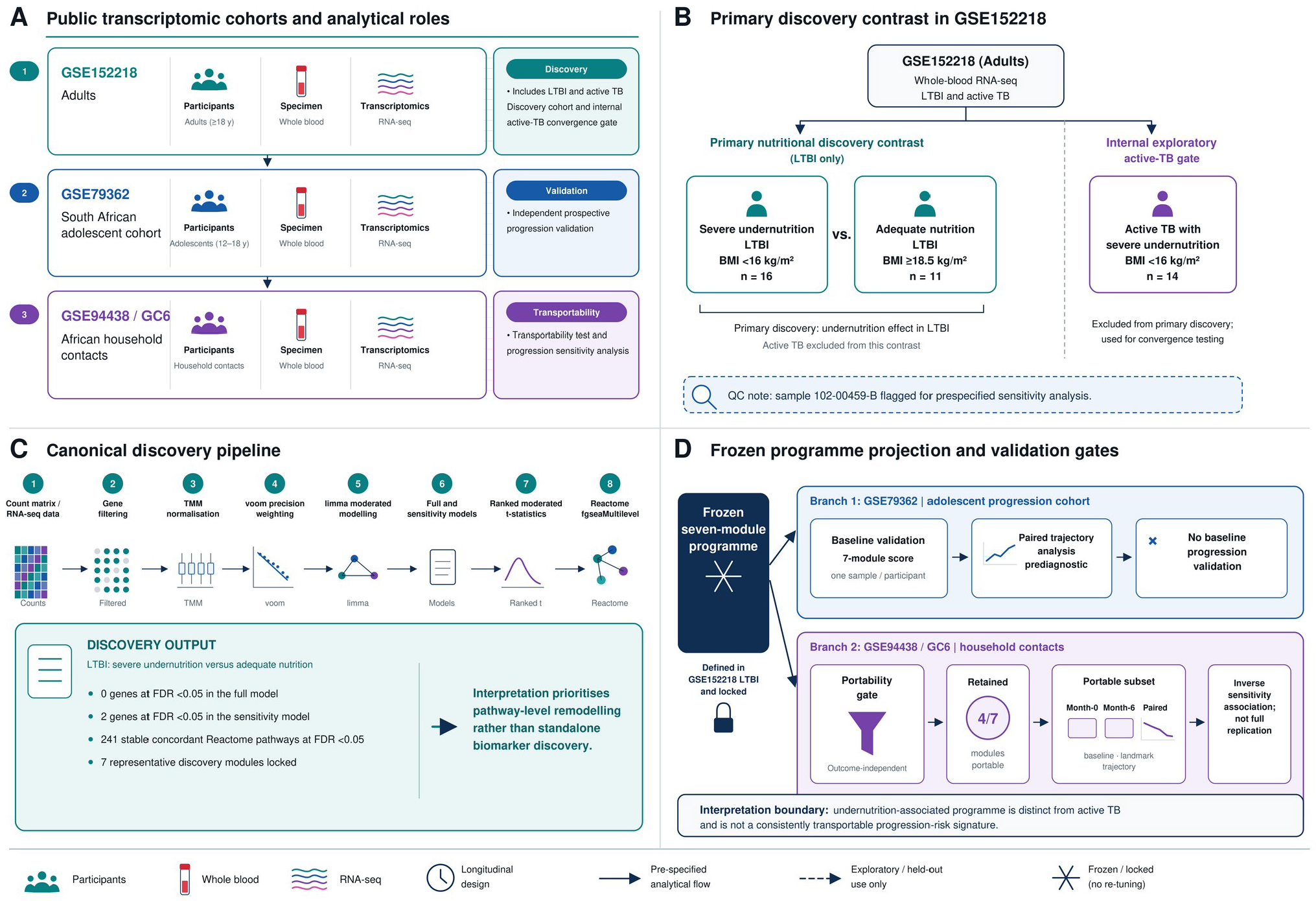
Study design and analytical framework. (A), Public transcriptomic cohorts and their roles: discovery in GSE152218, comparison with active TB within GSE152218, prospective testing in GSE79362, and household-contact testing in Grand Challenges 6-74 (GC6-74; GSE94438). (B), Primary discovery comparison restricted to adults with LTBI and severe undernutrition versus adequate nutritional status. (C), edgeR/limma-voom and Reactome analysis workflow. (D), Seven-module score tested in GSE79362 and GC6 after checking whether each predefined pathway module had sufficient assay representation. *Alt text: Four-panel schematic showing the three public transcriptomic cohorts and their analytical roles, the GSE152218 severe-undernutrition group comparison, the edgeR/limma-voom and Reactome analysis workflow, and testing of the seven-module score in GSE79362 and Grand Challenges 6-74 (GC6-74) after pathway-representation assessment*.

Severe undernutrition was defined as BMI <16 kg/m2 and compared with BMI >=18.5 kg/m2; BMI 16.0-18.49 kg/m2 was outside the prespecified discovery contrast. The analysis therefore estimates an extreme-phenotype difference rather than a graded BMI relationship.

### 2.2. Discovery comparison in GSE152218

The primary discovery comparison included adults with LTBI and available age, sex, and BMI data. In the source study, LTBI household contacts contributing to RNA sequencing were tuberculin skin test positive (TST >=5 mm), monitored for active-TB symptoms for 24 months, and did not progress to active TB [12,15]. We retained deposited TB-status classifications. Severe undernutrition included 16 adults and adequate nutrition 11; active-TB participants were excluded. Sample 102-00459-B was retained in the full analysis but excluded in the prespecified sensitivity analysis after quality-control review.

Participant-level inclusion, nutritional-status classification and the discovery-cohort audit are reported in Supplementary Table S1 (Supplementary material 1).

### 2.3. RNA-seq preprocessing and differential-expression modeling

Blood collection, RNA extraction, library preparation, and sequencing were performed by the original investigators [12,15]. We reanalyzed the deposited count matrix. Ensembl version suffixes were removed; genes with counts >=10 in at least three samples were retained; libraries were TMM-normalized in edgeR [16,17]; and voom/limma models adjusted for age and sex [18,19], with empirical-Bayes moderation and Benjamini-Hochberg FDR correction.

The full model included 27 adults; the sensitivity model excluded 102-00459-B and repeated filtering and normalization in 26 samples. Robustness was assessed by Spearman correlation of log2 fold-changes and directional agreement. Because this was a secondary reanalysis, sample size reflected eligible public records rather than prospective enrollment.

Complete discovery-model and sensitivity-analysis summary statistics are provided in Supplementary Table S2 (Supplementary material 1) and Supplementary Data S1 (Supplementary material 3).

### 2.4. Reactome pathway enrichment and seven-module summary

Pathway enrichment used fgseaMultilevel on limma moderated t-statistics [20]. Ensembl identifiers were mapped to HGNC symbols using GENCODE v49 [22]; duplicate symbols were resolved by retaining the statistic with greatest absolute moderated t value. Reactome gene sets were obtained from the Reactome knowledgebase [23], and pathways represented by 15–500 mapped genes were tested.

Pathways were stable if directionally concordant and FDR <0.05 in both models. Seven representative pathways summarized major immune and metabolic themes, with scoring directions fixed from the sensitivity model before external outcome testing. A higher module score indicated greater similarity to the severe-undernutrition pattern. These pathway scores are inferred from RNA expression, not direct protein or metabolite measurements.

The seven selected modules, enrichment statistics, and scoring directions are provided in Supplementary Table S3 (Supplementary material 1).

### 2.5. Comparison with active TB among severely undernourished adults

To test resemblance to active disease, we compared 14 adults with active TB and severe undernutrition with the 16 adults with LTBI and severe undernutrition in GSE152218. The same age- and sex-adjusted framework and outlier-excluded sensitivity analysis were used. Because the groups differed in age and sex and active TB can itself contribute to wasting, this comparison was exploratory and cannot establish whether undernutrition preceded or resulted from active disease.

Exact gene- and pathway-level comparison metrics are reported in Supplementary Table S4 (Supplementary material 1).

### 2.6. Prospective testing in GSE79362

GSE79362 is a longitudinal adolescent cohort in which participants without active TB at baseline were followed for incident TB [9,13]. Its curated expression matrix was accessed through curatedTBData [21]. The seven-module score was calculated after gene-wise standardization based on baseline samples. The primary analysis retained one baseline sample per participant and compared future progressors with non-progressors. Secondary analyses considered progression within 12 months and paired farthest-versus-nearest prediagnostic observations. Logistic regression adjusted for age, gender, ethnicity, and ACS cohort assignment.

Module representation and outcome analyses for GSE79362 are reported in Supplementary Table S5 (Supplementary material 1).

### 2.7. Prospective testing in Grand Challenges 6-74 (GC6-74; GSE94438)

GSE94438/GC6 is a longitudinal household-contact cohort followed for incident TB [10,14]. Its curated expression matrix was accessed through curatedTBData [21]. Before outcome testing, each predefined pathway was assessed for representation in GC6 relative to GSE79362. A module entered the primary GC6 analysis when GC6 contained at least 60% as many measurable pathway genes as GSE79362 and at least 20 genes. Because this criterion compares pathway gene counts rather than exact identity, post hoc robustness analyses rescored the four qualifying modules using genes measurable in both cohorts and applied a stricter identity-overlap rule (>=60% of GSE79362-represented genes shared with GC6 and >=20 shared genes). Analyses evaluated month-0 progression, 12-month progression, a month-6 landmark, and paired prediagnostic trajectories; baseline models adjusted for age, gender, and region.

GC6 pathway-representation decisions and primary outcome analyses are reported in Supplementary Table S6 (Supplementary material 1). The roles of the three cohorts and major analytical stages are summarized in Supplementary Table S7 (Supplementary material 2).

### 2.8. Software and reproducibility

Analyses used R 4.5.3 with edgeR, limma, fgsea, curatedTBData 2.6.0, GENCODE v49, and Reactome [16,18-23]. The current reproducibility release (v1.1.0), including the revision-specific GC6 overlap-robustness materials, is publicly available on GitHub and archived on Zenodo [33].

Source datasets are available through GEO under GSE152218, GSE79362, and GSE94438 [13-15].

## 3. Results

### 3.1. Undernutrition-associated transcriptional changes in LTBI

The discovery analysis compared 16 adults with LTBI and severe undernutrition with 11 adults with adequate nutrition. The primary limma-voom model tested 20,382 genes and identified none at FDR <0.05. The sensitivity model tested 20,059 genes; two reached FDR <0.05 and nine reached FDR <0.10. Because no gene met the primary FDR threshold, the two sensitivity-only findings were not interpreted as robust biomarkers. Effect estimates remained highly concordant (log2 fold-change Spearman rho=0.928; 90.7% directional agreement).

Pathway evidence was stronger. The primary model identified 311 pathways at FDR <0.05 and the sensitivity model 264; 241 were significant and directionally concordant in both, and 293 were concordant at FDR <0.10 (NES Spearman rho=0.954; 89.9% directional agreement). Stable pathways included higher complement activation, phospholipid/phagocytosis, B-cell regulation, and vascular interactions, with lower neutrophil degranulation, autophagy, and TNFR2 non-canonical NF-kB signaling. These are pathway-level RNA-expression findings rather than direct protein or metabolite measurements.

Figure 2 and Supplementary Tables S2 and S3 (Supplementary material 1) summarize discovery robustness and the seven representative pathways.

**Figure 2.**
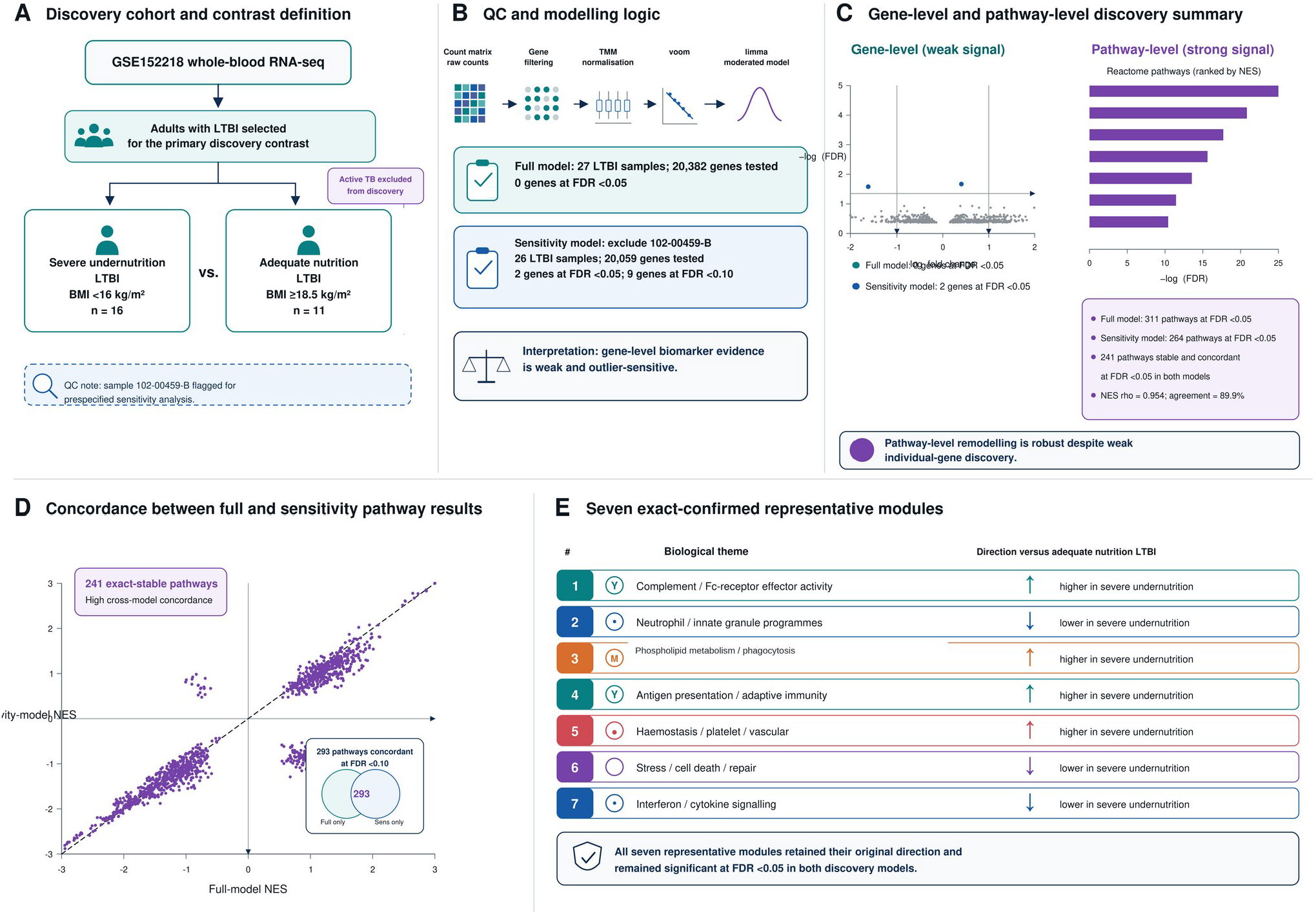
Severe-undernutrition-associated pathway changes among adults with LTBI. (A), Discovery group comparison in GSE152218. (B), Differential-expression workflow and prespecified sensitivity analysis. (C), Limited gene-level evidence versus robust Reactome pathway enrichment: 311 pathways reached FDR <0.05 in the full model, 264 in the sensitivity model, and 241 were significant and changed in the same direction in both models. (D), Concordance of Reactome normalized enrichment scores between models (Spearman rho=0.954; direction agreement=89.9%). (E), Seven representative immune and metabolic transcriptional pathways used in subsequent cohort tests. *Alt text: Five-panel summary of the discovery analysis showing cohort and quality-control logic, weak gene-level evidence, robust Reactome pathway enrichment, high concordance between full and sensitivity models, and the directions of seven representative immune and metabolic transcriptional pathways*.

### 3.2. Comparison with active TB among severely undernourished adults

Across 19,787 genes, the LTBI nutritional-status effect and the active-TB-versus-LTBI effect among severely undernourished adults were inversely associated (log2 fold-change rho=-0.399; moderated t rho=-0.427); only 34.9% of genes changed in the same direction.

The same pattern was evident across 1,274 Reactome pathways (NES rho=-0.390; 34.2% same direction). Of 241 stable undernutrition-associated pathways, 63 changed in the same direction and 178 in the opposite direction in active TB; 117 opposite-direction pathways were significant there. Thus, the severe-undernutrition LTBI pattern was predominantly unlike active-TB biology. Because disease-status groups differed in age and sex and active TB may contribute to wasting, this comparison remains exploratory.

Figure 3 summarizes the gene- and pathway-level comparison with active TB among severely undernourished adults.

**Figure 3.**
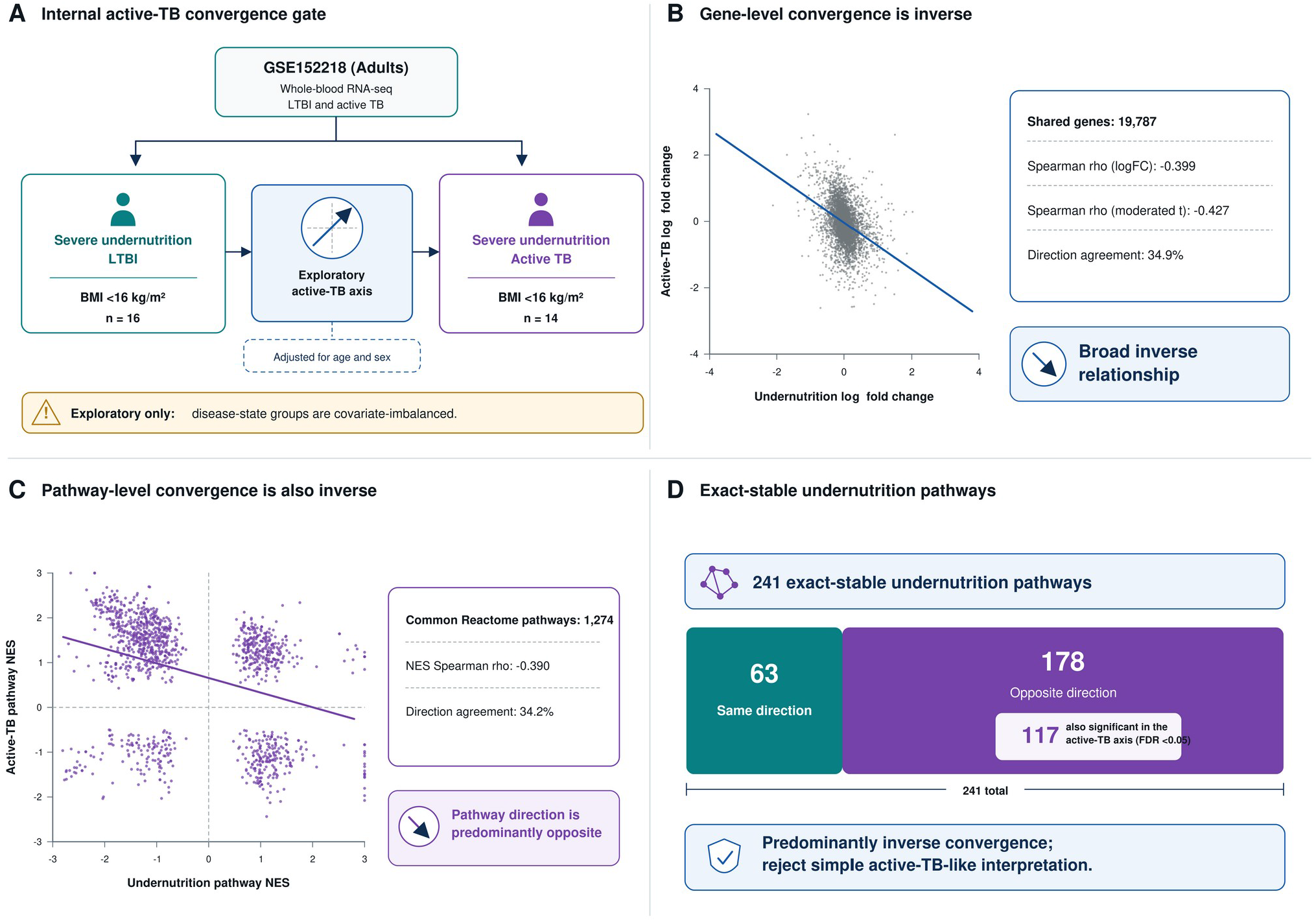
Comparison of undernutrition-associated transcriptional changes with active TB among severely undernourished adults. (A), Within-GSE152218 comparison of severe undernutrition versus adequate nutrition among adults with LTBI and active TB versus LTBI among severely undernourished adults, including the age- and sex-imbalance caution. (B), Gene-level comparison between the two sets of log2 fold-changes (Spearman rho=-0.399). (C), Reactome normalized enrichment score comparison (Spearman rho=-0.390). (D), Summary of the 241 stable undernutrition-associated pathways, showing 63 in the same direction and 178 in the opposite direction in the active-TB comparison. *Alt text: Four-panel comparison showing the two within-GSE152218 group comparisons, their gene-level effect correlation, their Reactome normalized enrichment score correlation, and counts of stable undernutrition-associated pathways that changed in the same versus opposite direction in active TB*.

### 3.3. Prospective testing in GSE79362

In GSE79362, the baseline analysis included 28 future progressors and 100 non-progressors. The seven-module score was not associated with progression (median difference=-0.0205; Mann-Whitney P=0.5056; AUC=0.459; adjusted OR per SD=1.090; P=0.6922).

A baseline analysis restricted to 12-month progression was also non-significant (median difference=-0.0227; P=0.8645). Among 19 progressors with paired prediagnostic samples, the score increased slightly nearer diagnosis (median change=0.0494; P=0.0671), without statistical significance.

Figure 4 summarizes the GSE79362 prospective analyses.

**Figure 4.**
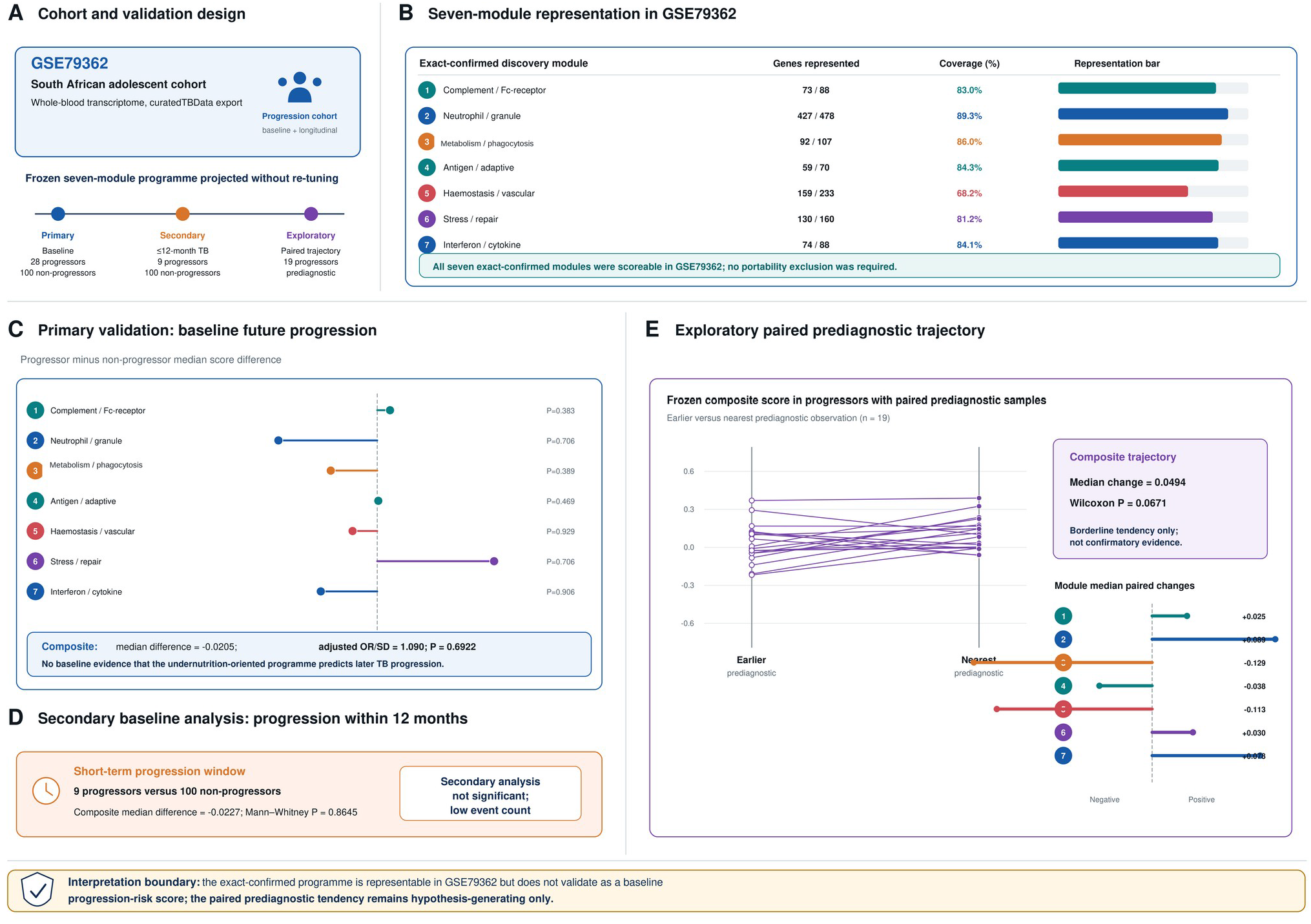
Testing the seven-module score for future TB in GSE79362. (A), Longitudinal adolescent-cohort design and baseline primary analysis. (B), Gene representation of the seven modules in the GSE79362 expression matrix. (C), Baseline seven-module scores in future progressors and non-progressors (adjusted OR per SD=1.090; P=0.6922). (D), Secondary analysis among participants who progressed within 12 months. (E), Paired prediagnostic trajectories among 19 progressors, showing a non-significant trend nearer diagnosis (P=0.0671). *Alt text: Five-panel evaluation in GSE79362 showing cohort design, module gene coverage, baseline combined-score comparison and adjusted association with progression, a 12-month progression analysis, and paired prediagnostic score trajectories*.

### 3.4. Prospective testing in GC6-74

In the second prospective cohort, GC6-74 (hereafter GC6), not all seven modules could be evaluated because three contained too few measured genes.

GC6 measured 5 of 73 complement/Fc-receptor genes, 23 of 92 phospholipid/phagocytosis genes, and 5 of 59 antigen/adaptive-immunity genes relative to GSE79362. Four modules met the prespecified representation criterion: neutrophil/innate granule activity, hemostasis/vascular activity, stress/cell-death/repair, and interferon/cytokine signaling. Representative undernutrition-associated pathways and their GC6 assay-representation status are detailed in Supplementary Table S8 (Supplementary material 2).

At month 0, 64 future progressors were compared with 208 non-progressors. The four-module score was lower among progressors (median difference=-0.0524; Mann-Whitney P=0.0220; AUC=0.405) and inversely associated with progression after adjustment for age, gender, and region (OR per SD=0.711; P=0.0218).

A post hoc gene-identity audit showed very high overlap for three qualifying modules; hemostasis/vascular activity shared 57.2% of its GSE79362-represented genes with GC6. The inverse baseline association was essentially unchanged when the four-module score used only shared genes (adjusted OR per SD=0.713; P=0.0224) and persisted after excluding hemostasis under the stricter identity-overlap rule (three-module OR=0.696; P=0.0145).

Secondary analyses were non-confirmatory: the 12-month baseline association did not reach significance (OR=0.710; P=0.0588), the month-6 landmark was null (OR=0.711; P=0.2851), and paired prediagnostic measurements among 20 progressors showed no consistent increase nearer diagnosis (median change=-0.0977; P=0.2549). Shared-gene and strict-overlap scores were likewise non-confirmatory. Thus, GC6 showed a robust inverse baseline association without consistent longitudinal confirmation.

Figure 5 summarizes the GC6 pathway-representation assessment and progression analyses. Supplementary Tables S9 and S10 (Supplementary material 2) summarize the major analytical questions, interpretation boundaries, and post hoc GC6 gene-identity overlap and score-sensitivity analyses; the corresponding machine-readable robustness outputs are provided in Supplementary material 4.

**Figure 5.**
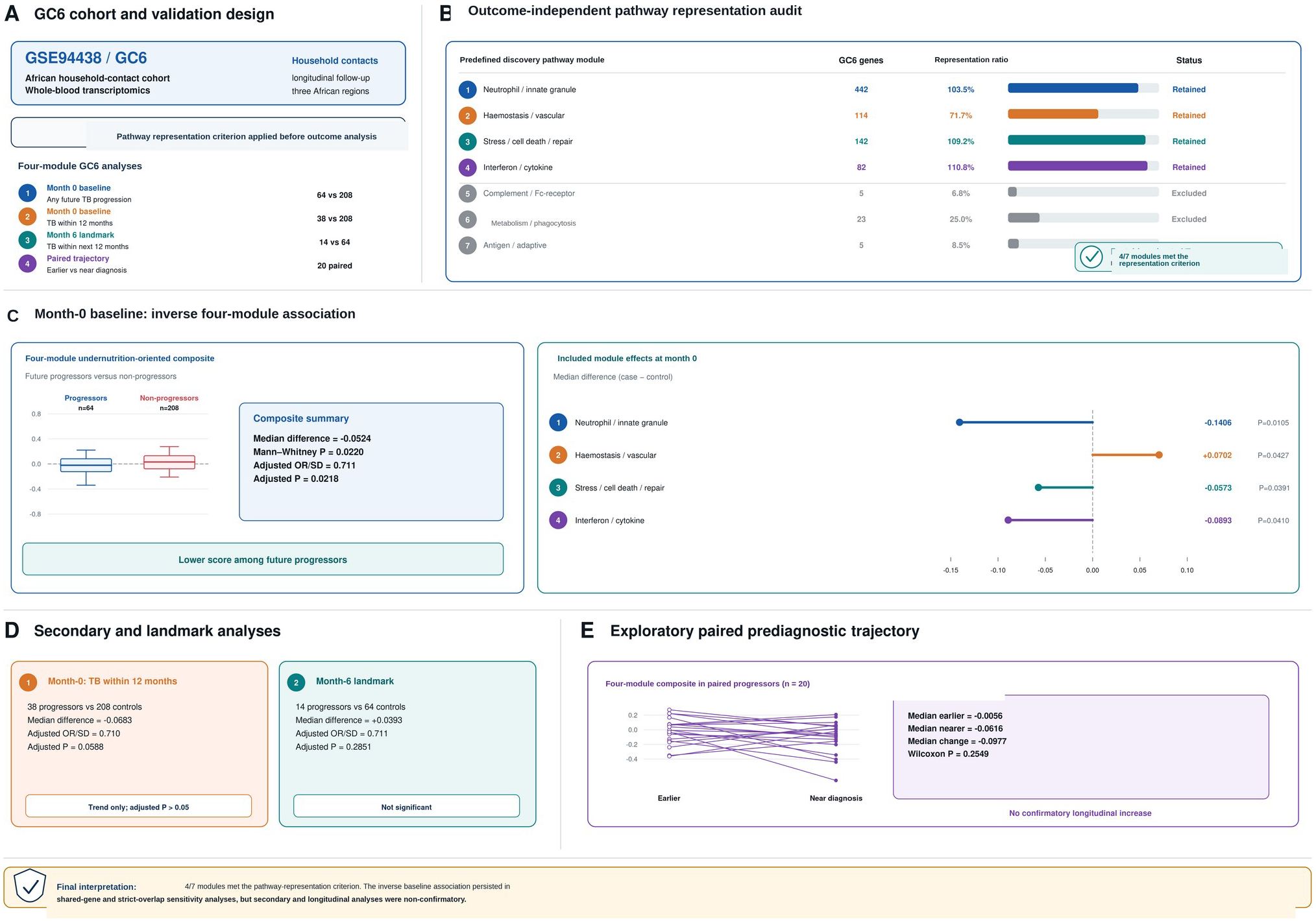
Testing the seven-module score in Grand Challenges 6-74 (GC6-74; GSE94438) after pathway-representation assessment. (A), Household-contact cohort design and the prespecified representation criterion applied before outcome analysis. (B), GC6:GSE79362 pathway gene-count ratios for the seven modules; four met the prespecified criterion. (C), Month-0 baseline scores for the four evaluable modules among future progressors and non-progressors (adjusted OR per SD=0.711; P=0.0218). The inverse baseline association was preserved in post hoc shared-gene and stricter identity-overlap sensitivity analyses (Supplementary Table S10). (D), Secondary landmark analyses. (E), Paired prediagnostic analysis among 20 progressors, which did not provide confirmatory longitudinal evidence. *Alt text: Five-panel GC6-74 analysis showing household-contact design, pathway-representation assessment, the four-module baseline score, secondary landmark analyses, paired prediagnostic trajectories, and a note that shared-gene sensitivity analyses preserved the inverse baseline association*.

## 4. Discussion

Severe undernutrition during LTBI was associated with reproducible immune- and metabolism-related transcriptional remodeling despite no primary-model FDR-significant genes. This extends reports of altered inflammatory and TB-risk-signature activity in malnourished LTBI [11,12] and attenuated M. tuberculosis-specific Th1 responses in undernourished contacts [32]. We further tested whether this nutritional transcriptional state resembled active TB or predicted progression; it did neither consistently across the analyses.

The pathway findings are biologically coherent but must be interpreted at the level measured. Higher enrichment involved complement/Fc-receptor activity, B-cell regulation, phospholipid/phagocytosis biology, and vascular interactions, whereas neutrophil degranulation, autophagy, and TNFR2 non-canonical NF-kB signaling were lower. Whole-blood RNA sequencing cannot establish corresponding protein, cytokine, metabolite, or flux changes, identify causal cell populations, or determine whether the patterns are protective, dysfunctional, or compensatory.

The predominantly inverse relationship with active TB argues against severe undernutrition during LTBI being simply an early active-disease transcriptional state. Active TB typically shows interferon-inducible and neutrophil-linked signatures [7,8], whereas the nutritional pattern showed lower neutrophil-degranulation and cytokine-signaling activity and broad opposite-direction pathway changes. Nutritional stress may therefore affect immune competence differently from established disease. This remains hypothesis-generating because disease-status groups differed in age and sex and active TB can itself cause wasting.

The external cohorts tested a nutritional-state score, not an incipient-TB classifier. In GSE79362 it was unrelated to baseline progression, and established progression signatures generally perform best nearer diagnosis [9,10,25-27]. These findings do not invalidate the undernutrition-associated biology; rather, they show that a transcriptional state defined by nutritional phenotype is not interchangeable with a short-horizon progression signature.

GC6 illustrates cross-study technical constraints. Three modules had insufficient pathway representation, preventing full seven-module evaluation. The four-module baseline score was inversely associated with progression; this association persisted with shared-gene scoring and after excluding hemostasis under a stricter identity-overlap rule. However, 12-month, month-6, and paired analyses were non-confirmatory. Because transcriptomic-signature performance varies with population, comparator, prediction horizon, and platform [28-30], the inverse baseline association should not be interpreted as protective undernutrition but as a cohort- and assay-dependent association.

Limitations include the modest discovery sample, an extreme BMI contrast (<16 vs >=18.5 kg/m2) that does not estimate a graded relationship, age/sex imbalance and possible reverse causality in the active-TB comparison, and external cohorts lacking the same severe-undernutrition phenotype. The external analyses therefore test whether the transcriptional score predicts future TB, not whether they replicate the nutritional effect itself. Finally, pathway enrichment is RNA-based and requires validation in prospectively nutrition-phenotyped cohorts with cell-resolved and orthogonal protein/metabolic measurements [31].

## 5. Conclusions

Severe undernutrition during LTBI is associated with reproducible immune- and metabolism-related transcriptional remodeling rather than a robust individual-gene signature. The pattern differs substantially from active-TB biology and does not consistently identify future TB progression across independent cohorts, supporting its interpretation as a distinct host state requiring prospective validation.

## Supporting information

GC6 Robustness Addendum

## Ethics statement

This secondary analysis used de-identified, publicly available datasets and involved no new recruitment, specimen collection, or intervention. Ethics approvals and consent procedures for the original studies are described in the source cohort publications [9,10,12]. Participant consent for publication is not applicable to the secondary reanalysis of publicly available, de-identified transcriptomic data.

## Data availability

Source transcriptomic datasets analysed in this study are publicly available through the Gene Expression Omnibus under GSE152218, GSE79362, and GSE94438 [13-15]. Curated external-cohort expression matrices were accessed through curatedTBData [21]. The reproducibility compendium, including analysis scripts, module definitions, derived outputs, metadata audits, session information, checksums, and GC6 overlap-robustness materials, is available at GitHub (https://github.com/rmaghembe1/tb-undernutrition-transcriptomics) and archived on Zenodo as v1.1.0 (https://doi.org/10.5281/zenodo.23119331) [33]. Supporting machine-readable outputs are also provided in Supplementary materials 3 and 4.

## Acknowledgements

The authors thank the participants and investigators of the original studies and the public repositories and curatedTBData resource that enabled this secondary analysis.

## CRediT authorship contribution statement

Conceptualization: RSM, SKH, SGM. Methodology: RSM. Formal analysis: RSM. Data curation: RSM. Validation: MN, RFM, SB, PS, WEJ. Supervision: SKH, SGM. Project administration: SKH, SGM. Funding acquisition: SKH, SGM. Writing - original draft: RSM. Writing - review and editing: RSM, VIK, PMM, RFM, SB, KMN, LN, MN, JMJ, WEJ, PS, SKH, SGM, SS. All authors read and approved the final manuscript.

## Declaration of generative AI and AI-assisted technologies in the writing process

During the preparation of this work, the authors used OpenAI ChatGPT (GPT-5.5 Thinking and GPT-5.6 Sol; OpenAI; accessed May-October 2026) for manuscript organization, language refinement, and journal-format alignment. The tool was not used to create or alter primary research data. The authors reviewed and edited all AI-assisted content, verified the scientific content and statistical values, and take full responsibility for the content of the publication.

## Supplementary material

Supplementary material 1 (.docx). Supplementary Methods and Tables S1-S6: detailed supplementary methods and Tables S1-S6 supporting cohort audits, discovery and sensitivity analyses, module definitions, active-TB comparison, GSE79362 testing, and GC6 primary analyses.

Supplementary material 2 (.docx). Supplementary Tables S7-S10: dataset roles, representative pathway and assay-representation summaries, interpretation boundaries, and the GC6 gene-identity overlap and score-sensitivity audit.

Supplementary material 3 (.zip). Supplementary Data S1: exact R outputs, analysis scripts, checksums, session and reproducibility materials supporting the reported analyses.

Supplementary material 4 (.zip). GC6 Robustness Addendum: shared-gene and strict identity-overlap code and machine-readable outputs for the GC6 robustness analyses.

