## Supplementary Methods and Tables S1-S6 for "Distinct transcriptional remodeling of immune and metabolic pathways in severe undernutrition during latent tuberculosis infection does not consistently predict tuberculosis progression across independent cohorts"

### Supplementary Material

*Supplementary Methods, Tables S1-S6 and Supplementary Data S1 manifest*

#### Supplementary Methods

##### Datasets and prespecified analytical structure

These supplementary resources report the exact R-confirmed outputs supporting the staged secondary analysis. GSE152218 provided the severe-undernutrition discovery contrast in adults with latent tuberculosis infection (LTBI) and the exploratory internal active-TB gate. GSE79362 provided prospective programme projection, and GSE94438/GC6 provided outcome-independent assay-portability-gated projection. The external cohorts test transportability of the discovery-defined programme rather than direct replication of the BMI-defined nutritional contrast.

##### Discovery cohort definition and sensitivity analysis

Adults with LTBI in GSE152218 were selected when BMI, age and sex metadata were available. Severe undernutrition was defined as BMI <16 kg/m2 and adequate nutritional status as BMI >=18.5 kg/m2. The complete model included 16 severely undernourished and 11 adequately nourished participants. Sample 102-00459-B was retained in the complete model and omitted in the prespecified sensitivity model following quality-control review.

##### Differential-expression modelling and Reactome enrichment

Gene-level modelling used trimmed-mean-of-M-values-normalised counts and limma-voom precision-weighted linear models adjusted for age and sex. Reactome enrichment used fgseaMultilevel on moderated t-statistic rankings after Ensembl-to-symbol harmonisation. A stable pathway required concordant direction and false discovery rate <0.05 in both complete and sensitivity models. Seven representative modules were locked before external outcome testing.

##### Internal active-TB convergence gate

The internal reference axis compared adults with severe undernutrition and active TB with adults with severe undernutrition and LTBI. Gene-level log2-fold-change and moderated-t convergence and pathway-level normalised enrichment-score convergence were assessed. This gate is interpreted as exploratory because of covariate imbalance between disease-state groups.

##### Prospective projection and assay portability

All seven exact-confirmed modules were scoreable in GSE79362. In GC6, an outcome-independent portability rule was applied before outcome testing: a module was retained only if the GC6 curated assay represented at least 60% of genes represented in GSE79362 and included at least 20 genes. Consequently, only four technically portable modules were evaluated in GC6 outcome analyses.

##### Reproducibility archive

Supplementary Data S1 supplies the exact machine-readable output tables, frozen module definition file, metadata audits, R scripts and session-information files supporting the numerical findings reported in the article and in Tables S1-S6.

#### Supplementary Tables

##### Table S1. Discovery cohort sample audit for the GSE152218 adult LTBI nutritional-status comparison.

| GEO sample | Participant ID | Age | Sex | BMI (kg/m2) | Severe-undernutrition group | Included |
|---|---|---|---|---|---|---|
| GSM4609386 | 102-00042-B | 45 | female | 14.81 | Severe undernutrition (<16) | Yes |
| GSM4609387 | 102-00088-B | 23 | male | 13.92 | Severe undernutrition (<16) | Yes |

| GEO sample | Participant ID | Age | Sex | BMI (kg/m2) | Severe-undernutrition group | Included |
|---|---|---|---|---|---|---|
| GSM4609388 | 102-00129-B0 | 25 | male | 24.15 | Not severe (>=16) | Yes |
| GSM4609389 | 102-00131-B | 40 | female | 15.09 | Severe undernutrition (<16) | Yes |
| GSM4609390 | 102-00171-B | 39 | female | 28.12 | Not severe (>=16) | Yes |
| GSM4609391 | 102-00188-B | 21 | female | 22.87 | Not severe (>=16) | Yes |
| GSM4609393 | 102-00217-B | 40 | female | 27.81 | Not severe (>=16) | Yes |
| GSM4609395 | 102-00240-B | 50 | female | 27.44 | Not severe (>=16) | Yes |
| GSM4609396 | 102-00241-B | 19 | female | 13.87 | Severe undernutrition (<16) | Yes |
| GSM4609397 | 102-00246-B | 34 | female | 15.61 | Severe undernutrition (<16) | Yes |
| GSM4609400 | 102-00265-B0 | 24 | female | 14.21 | Severe undernutrition (<16) | Yes |
| GSM4609401 | 102-00275-B | 46 | male | 33.16 | Not severe (>=16) | Yes |
| GSM4609402 | 102-00277-B | 22 | male | 19.52 | Not severe (>=16) | Yes |
| GSM4609404 | 102-00291-B | 23 | female | 21.01 | Not severe (>=16) | Yes |
| GSM4609405 | 102-00298-B | 22 | female | 30.83 | Not severe (>=16) | Yes |
| GSM4609407 | 102-00304-B | 60 | male | 19.09 | Not severe (>=16) | Yes |
| GSM4609409 | 102-00308-B0 | 37 | female | 26.31 | Not severe (>=16) | Yes |
| GSM4609410 | 102-00310-B | 18 | male | 15.68 | Severe undernutrition (<16) | Yes |
| GSM4609413 | 102-00323-B | 20 | male | 15.72 | Severe undernutrition (<16) | Yes |
| GSM4609421 | 102-00350-B | 36 | female | 15.66 | Severe undernutrition (<16) | Yes |
| GSM4609423 | 102-00365-B | 46 | female | 14.38 | Severe undernutrition (<16) | Yes |
| GSM4609424 | 102-00373-B | 54 | male | 13.72 | Severe undernutrition (<16) | Yes |
| GSM4609426 | 102-00374-B0 | 22 | female | 14.04 | Severe undernutrition (<16) | Yes |
| GSM4609430 | 102-00459-B | 19 | female | 15.82 | Severe undernutrition (<16) | Yes |
| GSM4609431 | 102-00467-B | 57 | female | 15.81 | Severe undernutrition (<16) | Yes |
| GSM4609432 | 102-00525-B | 31 | male | 15.17 | Severe undernutrition (<16) | Yes |
| GSM4609433 | 102-00548-B | 30 | male | 15.81 | Severe undernutrition (<16) | Yes |

#### Table S2. Canonical discovery and sensitivity-analysis robustness summary.

| Output type | Metric/model | Samples | Genes tested | Value / FDR<0.05 | FDR<0.10 |
|---|---|---|---|---|---|
| Differential-expression model | all_27_samples | 27 | 20382 | 0 | 0 |
| Differential-expression model | sensitivity_excluding_102-00459-B | 26 | 20059 | 2 | 9 |
| Gene-level robustness | Spearman correlation of log2FC | - | - | 0.928305 | - |
| Gene-level robustness | Direction agreement | - | - | 0.906974 | - |
| Gene-level robustness | Shared genes at FDR <0.05 | - | - | 0 | - |
| Reactome robustness | Full model pathways at FDR <0.05 | - | - | 311 | - |
| Reactome robustness | Sensitivity model pathways at FDR <0.05 | - | - | 264 | - |
| Reactome robustness | Stable concordant pathways at FDR <0.05 | - | - | 241 | - |
| Reactome robustness | Stable concordant pathways at FDR <0.10 | - | - | 293 | - |
| Reactome robustness | Pathway NES Spearman correlation | - | - | 0.95446 | - |
| Reactome robustness | Pathway direction agreement | - | - | 0.898903 | - |

#### Table S3. Exact-confirmed seven-module programme used for prospective projection.

| Theme | Representative pathway | Locked direction | Full NES | Full FDR | Sensitivity NES | Sensitivity FDR |
|---|---|---|---|---|---|---|
| Complement / Fc-receptor effector activity | Classical antibody-mediated complement activation | Higher in severe undernutrition | 3.555 | 5.79e-31 | 3.567 | 2.57e-32 |
| Neutrophil / innate granule programmes | Neutrophil degranulation | Lower in severe undernutrition | -2.788 | 3.03e-32 | -2.654 | 1.41e-28 |
| Metabolism / mitochondrial energetics | Role of phospholipids in phagocytosis | Higher in severe undernutrition | 3.413 | 2.30e-26 | 3.336 | 5.80e-28 |
| Antigen presentation / adaptive immunity | CD22 mediated BCR regulation | Higher in severe undernutrition | 3.485 | 4.21e-27 | 3.488 | 2.48e-28 |
| Haemostasis / platelet / vascular | Cell surface interactions at the vascular wall | Higher in severe undernutrition | 2.864 | 1.78e-21 | 2.830 | 4.45e-19 |
| Stress / cell death / repair | Autophagy | Lower in severe undernutrition | -2.087 | 3.05e-06 | -2.171 | 1.90e-07 |
| Interferon / cytokine signalling | TNFR2 non-canonical NF-kB pathway | Lower in severe undernutrition | -1.953 | 9.54e-04 | -1.907 | 4.72e-04 |

#### Table S4. Internal active-TB convergence-gate summary.

| Evidence level | Metric | Value |
|---|---|---|
| Gene-level | Shared genes | 19787 |
| Gene-level | Spearman correlation of logFC | -0.398962 |
| Gene-level | Spearman correlation of moderated t | -0.42693 |
| Gene-level | Directional agreement | 0.349371 |
| Pathway-level | Common Reactome pathways | 1274 |
| Pathway-level | Pathway NES Spearman correlation | -0.390255 |
| Pathway-level | Pathway direction agreement | 0.342229 |
| Pathway-level | Exact stable undernutrition pathways evaluated | 241 |
| Pathway-level | Exact stable pathways same direction | 63 |
| Pathway-level | Exact stable pathways opposite direction | 178 |
| Pathway-level | Opposite-direction stable pathways with activeTB FDR <0.05 | 117 |

*Interpretation boundary: this comparison is an exploratory mechanistic gate rather than independent clinical validation.*

#### Table S5. GSE79362 programme representation and prospective projection outcomes.

| Module theme | Represented genes | Coverage | Projection status |
|---|---|---|---|
| Complement / Fc-receptor effector activity | 73 | 83.0% | Scoreable |
| Neutrophil / innate granule programmes | 427 | 89.3% | Scoreable |
| Metabolism / mitochondrial energetics | 92 | 86.0% | Scoreable |
| Antigen presentation / adaptive immunity | 59 | 84.3% | Scoreable |
| Haemostasis / platelet / vascular | 159 | 68.2% | Scoreable |
| Stress / cell death / repair | 130 | 81.2% | Scoreable |
| Interferon / cytokine signalling | 74 | 84.1% | Scoreable |

| Analysis | Progressors/paired n | Controls | Median difference/change | Unadjusted P | Adjusted OR/SD | Adjusted P |
|---|---|---|---|---|---|---|
| Baseline: future progression at any follow-up | 28 | 100 | -0.0205 | 0.5056 | 1.090 | 0.6922 |
| Baseline: TB progression within 12 months | 9 | 100 | -0.0227 | 0.8645 | - | - |
| Paired prediagnostic trajectory | 19 | - | 0.0494 | 0.0671 | - | - |

**Table S6. GC6 assay-portability gate and technically gated outcome analyses.**

| Module theme | GSE79362 genes | GC6 genes | Retention | Portability decision |
|---|---|---|---|---|
| Complement / Fc-receptor effector activity | 73 | 5 | 6.8% | Excluded |
| Neutrophil / innate granule programmes | 427 | 442 | 103.5% | Retained |
| Metabolism / mitochondrial energetics | 92 | 23 | 25.0% | Excluded |
| Antigen presentation / adaptive immunity | 59 | 5 | 8.5% | Excluded |
| Haemostasis / platelet / vascular | 159 | 114 | 71.7% | Retained |
| Stress / cell death / repair | 130 | 142 | 109.2% | Retained |
| Interferon / cytokine signalling | 74 | 82 | 110.8% | Retained |

| Analysis | Progressors/paired n | Controls | Median difference/change | Unadjusted P | Adjusted OR/SD | Adjusted P |
|---|---|---|---|---|---|---|
| Month 0 baseline: any future TB progression | 64 | 208 | -0.0524 | 0.0220 | 0.711 | 0.0218 |
| Month 0 baseline: progression within 12 months | 38 | 208 | -0.0683 | 0.0325 | 0.710 | 0.0588 |
| Month 6 landmark: TB within next 12 months | 14 | 64 | 0.0393 | 0.5363 | 0.711 | 0.2851 |
| Paired prediagnostic trajectory | 20 | - | -0.0977 | 0.2549 | - | - |

#### Supplementary Data S1. Exact-output and reproducibility archive

Supplementary Data S1 is provided as a ZIP archive containing exact R-confirmed output tables, frozen module definitions, metadata audit files, R scripts and R session-information files. A SHA-256 checksum manifest is included within the archive.
