## Supplementary Tables S7-S10 for "Distinct transcriptional remodeling of immune and metabolic pathways in severe undernutrition during latent tuberculosis infection does not consistently predict tuberculosis progression across independent cohorts"

### Distinct transcriptional remodeling of immune and metabolic pathways in severe undernutrition during latent tuberculosis infection does not consistently predict tuberculosis progression across independent cohorts

#### Supplementary Table S7. Public datasets, study populations and prespecified analytical roles.

| Dataset | Population and setting | Analytical use | Key design safeguard |
|---|---|---|---|
| GSE152218 | Adults with LTBI or active TB; India; whole-blood RNA-seq | Discovery contrast and exploratory active-TB comparison | Nutrition discovery restricted to adult LTBI; active-TB comparison treated as exploratory |
| GSE79362 | South African adolescents with M. tuberculosis infection followed longitudinally | Prospective seven-module score testing | Baseline primary analysis; paired trajectory secondary analysis |
| GSE94438 / GC6 | African household contacts sampled longitudinally after exposure | Progression testing after pathway-representation assessment | Outcome-independent pathway-representation criterion before scoring |

#### Supplementary Table S8. Representative undernutrition-associated pathways and GC6 assay-representation status.

| Biological theme | Representative Reactome pathway | Direction in severe undernutrition | Sensitivity NES | Sensitivity FDR | GC6 representation criterion |
|---|---|---|---|---|---|
| Complement / Fc-receptor effector activity | Classical antibody-mediated complement activation | Higher in severe undernutrition | 3.567 | 2.57e-32 | Did not meet |
| Neutrophil / innate granule programs | Neutrophil degranulation | Lower in severe undernutrition | -2.654 | 1.41e-28 | Met |
| Phospholipid metabolism / phagocytosis | Role of phospholipids in phagocytosis | Higher in severe undernutrition | 3.336 | 5.80e-28 | Did not meet |
| Antigen presentation / adaptive immunity | CD22 mediated BCR regulation | Higher in severe undernutrition | 3.488 | 2.48e-28 | Did not meet |
| Hemostasis / platelet / vascular | Cell surface interactions at the vascular wall | Higher in severe undernutrition | 2.830 | 4.45e-19 | Met |
| Stress / cell death / repair | Autophagy | Lower in severe undernutrition | -2.171 | 1.90e-07 | Met |
| Interferon / cytokine signaling | TNFR2 non-canonical NF-kB pathway | Lower in severe undernutrition | -1.907 | 4.72e-04 | Met |

#### Supplementary Table S9. Major analytical questions and interpretation boundaries.

| Analytical question | Exact result | Interpretation |
|---|---|---|
| Is severe undernutrition associated with differences in LTBI blood transcriptional biology? | 241 stable Reactome pathways; 7/7 representative pathways directionally confirmed in both models | Yes, at the coordinated pathway level |
| Does it yield a robust standalone gene biomarker signature? | 0 FDR-significant genes in the full discovery model | No |
| Does the undernutrition-associated pattern resemble active TB? | Pathway NES rho=-0.390; 178/241 stable pathways opposite | No; predominantly inverse in the exploratory internal comparison |
| Does it predict progression in GSE79362 at baseline? | Adjusted OR per SD=1.090; P=0.6922 | No evidence |
| Can all seven predefined modules be evaluated in GC6 under the prespecified representation criterion? | 4/7 modules meet the pathway-representation criterion | No; the full seven-module score cannot be evaluated in GC6 |
| Do the four evaluable GC6 modules associate with progression at baseline? | Adjusted OR per SD=0.711; P=0.0218; robust to shared-gene and strict-overlap sensitivity analyses | Inverse baseline association; not consistently confirmed in secondary or longitudinal analyses |

Supplementary Table S10. Post hoc GC6 gene-identity overlap audit and score-sensitivity analyses.

#### A. Gene-identity overlap for the seven predefined pathway modules.

| Module | GSE79362 genes | GC6 genes | Shared genes | Shared / GSE79362 (%) | Jaccard (%) | Prespecified count-ratio criterion | Strict identity-overlap criterion |
|---|---|---|---|---|---|---|---|
| Complement / Fc-receptor effector activity | 73 | 5 | 4 | 5.5 | 5.4 | Did not meet | Did not meet |
| Neutrophil / innate granule | 427 | 442 | 421 | 98.6 | 94.0 | Met | Met |
| Phospholipid metabolism / phagocytosis | 92 | 23 | 23 | 25.0 | 25.0 | Did not meet | Did not meet |
| Antigen presentation / adaptive immunity | 59 | 5 | 5 | 8.5 | 8.5 | Did not meet | Did not meet |
| Hemostasis / platelet / vascular | 159 | 114 | 91 | 57.2 | 50.0 | Met | Did not meet |
| Stress / cell death / repair | 130 | 142 | 130 | 100.0 | 91.5 | Met | Met |
| Interferon / cytokine signaling | 74 | 82 | 72 | 97.3 | 85.7 | Met | Met |

#### B. GC6 outcome sensitivity across alternative cross-assay score definitions.

| Analysis | Score definition | Cases / controls | Adjusted OR per SD | Adjusted P | Median difference / paired delta | Unadjusted P |
|---|---|---|---|---|---|---|
| Month 0: any future TB | Current 4-module | 64 / 208 | 0.711 | 0.02178 | -0.0524 | 0.02199 |
| Month 0: any future TB | Shared-gene 4-module | 64 / 208 | 0.713 | 0.02236 | -0.0553 | 0.01988 |
| Month 0: any future TB | Strict-overlap 3-module | 64 / 208 | 0.696 | 0.01445 | -0.0768 | 0.01508 |
| Month 0: TB within 12 months | Current 4-module | 38 / 208 | 0.710 | 0.05884 | -0.0683 | 0.03249 |
| Month 0: TB within 12 months | Shared-gene 4-module | 38 / 208 | 0.707 | 0.05454 | -0.0743 | 0.03034 |
| Month 0: TB within 12 months | Strict-overlap 3-module | 38 / 208 | 0.708 | 0.05416 | -0.0856 | 0.03310 |
| Month 6 landmark | Current 4-module | 14 / 64 | 0.711 | 0.28510 | +0.0393 | 0.53630 |
| Month 6 landmark | Shared-gene 4-module | 14 / 64 | 0.682 | 0.23010 | +0.0370 | 0.43090 |
| Month 6 landmark | Strict-overlap 3-module | 14 / 64 | 0.600 | 0.11760 | +0.0101 | 0.37950 |
| Paired prediagnostic | Current 4-module | 20 paired | - | - | -0.0977 | 0.2549 |
| Paired prediagnostic | Shared-gene 4-module | 20 paired | - | - | -0.1017 | 0.1978 |
| Paired prediagnostic | Strict-overlap 3-module | 20 paired | - | - | -0.1196 | 0.2250 |

Notes: The prespecified GC6 criterion compared pathway gene counts across assays: GC6 had to contain at least 60% as many measurable pathway genes as GSE79362 and at least 20 pathway genes. The shared-gene four-module sensitivity recalculated the same four modules using only genes measurable in both datasets. The strict-overlap three-module sensitivity additionally required at least 60% of the GSE79362-represented genes to be shared with GC6 and at least 20 shared genes; this excluded the hemostasis/vascular module. These analyses were post hoc robustness checks and do not replace the prespecified four-module analysis.
