## Supplementary material for "Distinct transcriptional remodeling of immune and metabolic pathways in severe undernutrition during latent tuberculosis infection does not consistently predict tuberculosis progression across independent cohorts": GC6 Robustness Addendum: 06_GC6_shared_gene_overlap_sensitivity.R

```
#!/usr/bin/env Rscript
# Revision-specific post hoc GC6 cross-assay overlap robustness analysis.
# Usage: Rscript 06_GC6_shared_gene_overlap_sensitivity.R /path/to/tb_undernutrition_transcriptomics /output/dir
suppressPackageStartupMessages(library(data.table))
args <- commandArgs(trailingOnly=TRUE)
if (length(args) < 1) stop("Provide project root as first argument")
root <- normalizePath(args[1], mustWork=TRUE)
out_dir <- if (length(args) >= 2) args[2] else getwd()
dir.create(out_dir, recursive=TRUE, showWarnings=FALSE)
source(file.path(root,"scripts/R/phaseB_helpers.R"))
read_expr <- function(path) {
  x <- fread(cmd=paste("gzip -dc",shQuote(path)))
  genes <- x[[1]]; mat <- as.matrix(x[,-1,with=FALSE]); rownames(mat) <- genes
  rowsum(mat,group=rownames(mat),reorder=FALSE) / as.numeric(table(factor(rownames(mat),levels=unique(rownames(mat)))))
}
gc6_expr <- read_expr(file.path(root,"data/external_validation/GSE94438/curatedTBData/GSE94438_assay_curated.tsv.gz"))
g793_expr <- read_expr(file.path(root,"data/external_validation/GSE79362/curatedTBData/GSE79362_assay_curated.tsv.gz"))
meta <- fread(file.path(root,"data/external_validation/GSE94438/curatedTBData/GSE94438_colData.tsv"),data.table=FALSE)
rownames(meta) <- meta$sample_id; gc6_expr <- gc6_expr[,meta$sample_id,drop=FALSE]
meta$TimeFromExposure_months <- parse_numeric_time(meta$TimeFromExposure); meta$TimeToTB_months <- parse_numeric_time(meta$TimeToTB)
meta$progressor <- meta$Progression=="Positive"; meta$labelled <- meta$Progression %in% c("Positive","Negative")
meta$Gender <- factor(meta$Gender); meta$GeographicalRegion <- factor(meta$GeographicalRegion); meta$Age <- as.numeric(meta$Age)
meta$months_sample_to_TB <- meta$TimeToTB_months-meta$TimeFromExposure_months
meta$prediagnosis_case <- meta$progressor & !is.na(meta$months_sample_to_TB) & meta$months_sample_to_TB>0
mods <- fread(file.path(root,"data/modules/exact_confirmed_seven_module_programme.tsv")); sets <- read_gmt_simple(file.path(root,"data/pathways/ReactomePathways.gmt"))
audit <- rbindlist(lapply(seq_len(nrow(mods)),function(i){
  gs <- sets[[mods$representative_pathway[i]]]; a <- intersect(gs,rownames(g793_expr)); b <- intersect(gs,rownames(gc6_expr)); sh <- intersect(a,b); un <- union(a,b)
  data.table(i=i,theme=mods$theme[i],GSE79362_genes=length(a),GC6_genes=length(b),shared_genes=length(sh),shared_over_GSE79362_pct=100*length(sh)/max(length(a),1),shared_over_GC6_pct=100*length(sh)/max(length(b),1),Jaccard_pct=100*length(sh)/max(length(un),1),current_representation_ratio_pct=100*length(b)/max(length(a),1))
}))
audit[,current_gate:=ifelse(current_representation_ratio_pct>=60 & GC6_genes>=20,"PASS","FAIL")]
audit[,strict_overlap_gate:=ifelse(shared_over_GSE79362_pct>=60 & shared_genes>=20,"PASS","FAIL")]
fwrite(audit,file.path(out_dir,"GC6_module_identity_overlap_v27.tsv"),sep="\t")
current_idx <- audit[current_gate=="PASS",i]; strict_idx <- audit[strict_overlap_gate=="PASS",i]
ids <- meta$sample_id[meta$labelled]; z <- zscore_rows(gc6_expr[,ids,drop=FALSE],reference_columns=ids)
make_composite <- function(idx,shared_only=FALSE){
  tmp <- data.table(sample_id=ids)
  for(j in idx){ pathway <- mods$representative_pathway[j]; gs <- if(shared_only) Reduce(intersect,list(sets[[pathway]],rownames(g793_expr),rownames(z))) else intersect(sets[[pathway]],rownames(z)); tmp[[mods$theme[j]]] <- colMeans(z[gs,,drop=FALSE],na.rm=TRUE)*mods$orientation_multiplier[j] }
  rowMeans(as.data.frame(tmp[,..mods$theme[idx]]),na.rm=TRUE)
}
scores <- data.table(sample_id=ids,CURRENT_4_MODULE=make_composite(current_idx,FALSE),SHARED_GENE_4_MODULE=make_composite(current_idx,TRUE),STRICT_OVERLAP_3_MODULE=make_composite(strict_idx,TRUE))
dat <- merge(meta[meta$labelled,],scores,by="sample_id"); score_names <- names(scores)[-1]
test_design <- function(df,outcome_name,label){ y <- as.logical(df[[outcome_name]]); rbindlist(lapply(score_names,function(score){ cases <- df[[score]][y]; controls <- df[[score]][!y]; wt <- wilcox.test(cases,controls,exact=FALSE); auc <- as.numeric(wt$statistic)/(sum(y)*sum(!y)); d <- df[complete.cases(df[,c(score,outcome_name,"Age","Gender","GeographicalRegion")]),]; d$score_z <- as.numeric(scale(d[[score]])); fit <- glm(as.formula(paste(outcome_name,"~ score_z + Age + Gender + GeographicalRegion")),data=d,family=binomial()); data.table(analysis=label,score=score,case_n=sum(y),control_n=sum(!y),median_difference=median(cases)-median(controls),mann_whitney_p=wt$p.value,AUC=auc,adjusted_OR_per_SD=exp(coef(fit)["score_z"]),adjusted_p=summary(fit)$coefficients["score_z","Pr(>|z|)"]) })) }
baseline <- dat[dat$TimeFromExposure_months==0,]; baseline$case <- baseline$progressor
base12 <- baseline[!baseline$progressor | (baseline$TimeToTB_months>=1 & baseline$TimeToTB_months<=12),]; base12$case <- base12$progressor & base12$TimeToTB_months>=1 & base12$TimeToTB_months<=12
month6 <- dat[dat$TimeFromExposure_months==6 & (!dat$progressor | dat$TimeToTB_months>6),]; month6$case <- month6$progressor & month6$TimeToTB_months>=7 & month6$TimeToTB_months<=18
binary <- rbindlist(list(test_design(baseline,"case","Month 0 baseline: any future TB progression"),test_design(base12,"case","Month 0 baseline: progression within 12 months"),test_design(month6,"case","Month 6 landmark: TB within next 12 months")))
fwrite(binary,file.path(out_dir,"GC6_score_definition_sensitivity_v27.tsv"),sep="\t")
# paired trajectories
pred <- dat[dat$prediagnosis_case,]; pairs <- list()
for(pid in unique(pred$PatientID)){ d <- pred[pred$PatientID==pid,]; if(length(unique(d$TimeFromExposure_months))>=2){ early <- d[which.min(d$TimeFromExposure_months),]; near <- d[which.min(d$months_sample_to_TB),]; if(early$sample_id!=near$sample_id){ rec <- data.table(PatientID=pid); for(score in score_names) rec[[paste0(score,"__delta")]] <- near[[score]]-early[[score]]; pairs[[as.character(pid)]] <- rec } } }
pairs <- rbindlist(pairs); traj <- rbindlist(lapply(score_names,function(score){ delta <- pairs[[paste0(score,"__delta")]]; w <- wilcox.test(delta,mu=0,exact=FALSE); data.table(analysis="Paired prediagnostic",score=score,paired_progressors=nrow(pairs),median_delta_near_minus_early=median(delta),wilcoxon_p=w$p.value) }))
fwrite(traj,file.path(out_dir,"GC6_paired_score_sensitivity_v27.tsv"),sep="\t")
```
