## Supplementary material for "Distinct transcriptional remodeling of immune and metabolic pathways in severe undernutrition during latent tuberculosis infection does not consistently predict tuberculosis progression across independent cohorts": GC6 Robustness Addendum: README_GC6_Robustness_Addendum.md

# Supplementary Data S1 Addendum - GC6 overlap robustness (v27)

This addendum documents the post hoc cross-assay robustness analyses introduced after audit of the original GC6 pathway-representation rule. The prespecified four-module analysis remains primary.

## Key results
- Current four-module baseline: adjusted OR/SD 0.711, P=0.02178.
- Shared-gene four-module baseline: adjusted OR/SD 0.713, P=0.02236.
- Strict-overlap three-module baseline: adjusted OR/SD 0.696, P=0.01445.
- Twelve-month, month-6, and paired prediagnostic analyses were non-confirmatory under all score definitions.

The strict overlap rule requires at least 60% of the GSE79362-represented pathway genes to be shared with GC6 and at least 20 shared genes; this excludes the hemostasis/vascular module.

## Files
- `06_GC6_shared_gene_overlap_sensitivity.R`: reproducible analysis script using the existing project inputs and helper functions.
- `GC6_module_identity_overlap_v27.tsv`: module-level gene-identity audit.
- `GC6_score_definition_sensitivity_v27.tsv`: binary-outcome sensitivity results.
- `GC6_Robustness_Audit_v27.xlsx`: two-sheet convenience workbook.

The original public v1.0.0 release remains immutable; this addendum is revision-specific and is intended for the manuscript v27 submission and a future v1.1.0 public update.
