## Supplementary Data S1 for "Distinct transcriptional remodeling of immune and metabolic pathways in severe undernutrition during latent tuberculosis infection does not consistently predict tuberculosis progression across independent cohorts": 00_install_confirmation_packages.R

```
#!/usr/bin/env Rscript
options(repos = c(CRAN = "https://cloud.r-project.org"))
options(timeout = 3600)

if (!requireNamespace("BiocManager", quietly = TRUE)) {
  install.packages("BiocManager")
}

cat("R version:\n")
print(R.version.string)
cat("\nBioconductor version before installation:\n")
print(BiocManager::version())

bioc_pkgs <- c("edgeR", "limma", "fgsea")
cran_pkgs <- c("data.table")

for (pkg in bioc_pkgs) {
  if (!requireNamespace(pkg, quietly = TRUE)) {
    BiocManager::install(pkg, ask = FALSE, update = FALSE)
  }
}
for (pkg in cran_pkgs) {
  if (!requireNamespace(pkg, quietly = TRUE)) {
    install.packages(pkg)
  }
}

required <- c(bioc_pkgs, cran_pkgs)
missing <- required[!vapply(required, requireNamespace, logical(1), quietly = TRUE)]
if (length(missing) > 0) {
  stop("Required packages still missing after installation: ", paste(missing, collapse = ", "))
}

cat("\nRequired confirmation packages are available:\n")
for (pkg in required) {
  cat(pkg, " version ", as.character(packageVersion(pkg)), "\n", sep = "")
}
cat("\nBioconductor package validation:\n")
print(BiocManager::valid())
```
