## Supplementary Data S1 for "Distinct transcriptional remodeling of immune and metabolic pathways in severe undernutrition during latent tuberculosis infection does not consistently predict tuberculosis progression across independent cohorts": 01_confirm_GSE152218_limma_voom.R

```
#!/usr/bin/env Rscript
# Exact canonical confirmation of the adult-LTBI discovery contrast.
# Models: severe undernutrition versus adequate nutrition, adjusted for age and sex.
# Full model plus prespecified sensitivity model excluding QC-flagged sample 102-00459-B.

suppressPackageStartupMessages({
  library(edgeR)
  library(limma)
  library(data.table)
})

args <- commandArgs(trailingOnly = TRUE)
project_root <- if (length(args) >= 1) normalizePath(args[1], mustWork = FALSE) else normalizePath(".", mustWork = FALSE)

count_file <- file.path(project_root, "data/raw/GSE152218_features_combined.txt.gz")
meta_file  <- file.path(project_root, "data/metadata/metadata_gse152218_adult_ltbi_primary.tsv")
out_dir    <- file.path(project_root, "results/R_confirmation/limma_voom")
dir.create(out_dir, recursive = TRUE, showWarnings = FALSE)

required_files <- c(count_file, meta_file)
missing <- required_files[!file.exists(required_files)]
if (length(missing) > 0) {
  stop("Missing required input(s):\n", paste(missing, collapse = "\n"))
}

meta <- fread(meta_file, data.table = FALSE, check.names = FALSE)
required_meta <- c("sample_accession", "subject_title", "severe_group", "age_years", "sex")
if (!all(required_meta %in% colnames(meta))) {
  stop("Metadata file lacks required fields: ", paste(setdiff(required_meta, colnames(meta)), collapse = ", "))
}

raw <- fread(cmd = paste("gzip -dc", shQuote(count_file)), data.table = FALSE, check.names = FALSE)
gene_col <- colnames(raw)[1]
gene_ids <- sub("\\..*$", "", as.character(raw[[gene_col]]))
if (!all(meta$subject_title %in% colnames(raw))) {
  stop("Not all subject_title identifiers are present in the count matrix.")
}

counts <- as.matrix(raw[, meta$subject_title, drop = FALSE])
mode(counts) <- "numeric"
rownames(counts) <- gene_ids
colnames(counts) <- meta$sample_accession
rownames(meta) <- meta$sample_accession
meta <- meta[colnames(counts), , drop = FALSE]

meta$nutrition <- ifelse(
  grepl("^Severe", meta$severe_group),
  "Severe_Undernutrition", "Adequate"
)
meta$nutrition <- factor(meta$nutrition, levels = c("Adequate", "Severe_Undernutrition"))
meta$sex <- factor(meta$sex)
meta$age_z <- as.numeric(scale(as.numeric(meta$age_years)))

run_model <- function(sample_keep, suffix) {
  model_meta <- meta[sample_keep, , drop = FALSE]
  model_counts <- counts[, rownames(model_meta), drop = FALSE]

  y <- DGEList(counts = model_counts)
  keep_gene <- rowSums(y$counts >= 10) >= 3
  y <- y[keep_gene, , keep.lib.sizes = FALSE]
  y <- calcNormFactors(y, method = "TMM")

  design <- model.matrix(~ nutrition + age_z + sex, data = model_meta)

  pdf(file.path(out_dir, paste0("voom_mean_variance_", suffix, ".pdf")), width = 7, height = 5.5)
  v <- voom(y, design, plot = TRUE)
  dev.off()

  fit <- lmFit(v, design)
  fit <- eBayes(fit)
  coef_name <- "nutritionSevere_Undernutrition"
  if (!coef_name %in% colnames(fit$coefficients)) {
    stop("Expected contrast coefficient not available: ", coef_name)
  }

  tab <- topTable(fit, coef = coef_name, number = Inf, sort.by = "P")
  tab$ensembl_gene_id <- rownames(tab)
  tab$direction <- ifelse(tab$logFC > 0, "Higher in severe undernutrition", "Lower in severe undernutrition")
  tab <- tab[, c("ensembl_gene_id", "logFC", "AveExpr", "t", "P.Value", "adj.P.Val", "B", "direction")]
  fwrite(tab, file.path(out_dir, paste0("limma_voom_", suffix, ".tsv")), sep = "\t")

  ranking <- tab[, c("ensembl_gene_id", "t")]
  ranking <- ranking[order(ranking$t, decreasing = TRUE), ]
  fwrite(ranking, file.path(out_dir, paste0("ranked_moderated_t_", suffix, ".rnk")), sep = "\t", col.names = FALSE)

  data.frame(
    model = suffix,
    samples = ncol(y),
    genes_tested = nrow(y),
    FDR_lt_0.05 = sum(tab$adj.P.Val < 0.05, na.rm = TRUE),
    FDR_lt_0.10 = sum(tab$adj.P.Val < 0.10, na.rm = TRUE),
    stringsAsFactors = FALSE
  )
}

full_keep <- rep(TRUE, nrow(meta))
sens_keep <- meta$subject_title != "102-00459-B"

summary_full <- run_model(full_keep, "all_27_samples")
summary_sens <- run_model(sens_keep, "sensitivity_excluding_102-00459-B")
summary_tab <- rbind(summary_full, summary_sens)
fwrite(summary_tab, file.path(out_dir, "limma_voom_model_summary.tsv"), sep = "\t")

full <- fread(file.path(out_dir, "limma_voom_all_27_samples.tsv"))
sens <- fread(file.path(out_dir, "limma_voom_sensitivity_excluding_102-00459-B.tsv"))
merged <- merge(full[, .(ensembl_gene_id, logFC_full = logFC, FDR_full = adj.P.Val)],
                sens[, .(ensembl_gene_id, logFC_sensitivity = logFC, FDR_sensitivity = adj.P.Val)],
                by = "ensembl_gene_id")
rho <- cor(merged$logFC_full, merged$logFC_sensitivity, method = "spearman", use = "complete.obs")
direction_agreement <- mean(sign(merged$logFC_full) == sign(merged$logFC_sensitivity), na.rm = TRUE)
shared_sig <- sum(merged$FDR_full < 0.05 & merged$FDR_sensitivity < 0.05, na.rm = TRUE)

robustness <- data.frame(
  metric = c("Spearman correlation of log2FC", "Direction agreement", "Shared genes at FDR <0.05"),
  value = c(rho, direction_agreement, shared_sig)
)
fwrite(robustness, file.path(out_dir, "limma_voom_robustness_summary.tsv"), sep = "\t")

writeLines(capture.output(sessionInfo()), file.path(out_dir, "sessionInfo_limma_voom.txt"))

cat("\nExact limma-voom confirmation complete.\n")
print(summary_tab)
cat("\nRobustness summary:\n")
print(robustness)
cat("\nResults directory:", out_dir, "\n")
```
