## Supplementary Data S1 for "Distinct transcriptional remodeling of immune and metabolic pathways in severe undernutrition during latent tuberculosis infection does not consistently predict tuberculosis progression across independent cohorts": 02_confirm_GSE152218_Reactome_fgsea.R

```
#!/usr/bin/env Rscript
# Exact Reactome preranked enrichment confirmation using fgseaMultilevel.
# Uses canonical limma-voom moderated t-statistics generated by script 01.

suppressPackageStartupMessages({
  library(fgsea)
  library(data.table)
})

args <- commandArgs(trailingOnly = TRUE)
project_root <- if (length(args) >= 1) normalizePath(args[1], mustWork = FALSE) else normalizePath(".", mustWork = FALSE)

limma_dir <- file.path(project_root, "results/R_confirmation/limma_voom")
out_dir   <- file.path(project_root, "results/R_confirmation/reactome_fgsea")
gmt_zip   <- file.path(project_root, "data/pathways/ReactomePathways.gmt.zip")
gmt_txt   <- file.path(project_root, "data/pathways/ReactomePathways.gmt")
map_file  <- file.path(project_root, "data/mapping/gencode.v49.ENSG_to_symbol.tsv.gz")

dir.create(out_dir, recursive = TRUE, showWarnings = FALSE)

if (!file.exists(gmt_txt)) {
  if (!file.exists(gmt_zip)) stop("Missing Reactome GMT ZIP or extracted GMT file.")
  unzip(gmt_zip, exdir = file.path(project_root, "data/pathways"))
}
if (!file.exists(gmt_txt)) stop("ReactomePathways.gmt was not found after ZIP extraction.")
if (!file.exists(map_file)) stop("Missing GENCODE Ensembl-to-symbol mapping file: ", map_file)

full_file <- file.path(limma_dir, "limma_voom_all_27_samples.tsv")
sens_file <- file.path(limma_dir, "limma_voom_sensitivity_excluding_102-00459-B.tsv")
if (!file.exists(full_file) || !file.exists(sens_file)) {
  stop("limma-voom outputs are missing. Run 01_confirm_GSE152218_limma_voom.R first.")
}

pathways <- gmtPathways(gmt_txt)
mapping <- fread(cmd = paste("gzip -dc", shQuote(map_file)))
mapping[, ensembl_gene_id := sub("\\..*$", "", ensembl_gene_id)]
mapping <- unique(mapping[!is.na(gene_symbol) & gene_symbol != "", .(ensembl_gene_id, gene_symbol)])

prepare_stats <- function(result_file, label) {
  tab <- fread(result_file)
  tab[, ensembl_gene_id := sub("\\..*$", "", ensembl_gene_id)]
  mapped <- merge(tab, mapping, by = "ensembl_gene_id", all = FALSE)
  mapped <- mapped[!is.na(t) & !is.na(gene_symbol) & gene_symbol != ""]
  mapped[, abs_t__ := abs(t)]
  setorder(mapped, -abs_t__)
  mapped[, abs_t__ := NULL]
  duplicates_removed <- nrow(mapped) - uniqueN(mapped$gene_symbol)
  mapped <- mapped[!duplicated(gene_symbol)]
  stats <- mapped$t
  names(stats) <- mapped$gene_symbol
  stats <- sort(stats, decreasing = TRUE)

  audit <- data.frame(
    model = label,
    limma_genes = nrow(tab),
    mapped_rows = nrow(merge(tab, mapping, by = "ensembl_gene_id", all = FALSE)),
    unique_symbols_used = length(stats),
    duplicate_symbols_removed = duplicates_removed,
    stringsAsFactors = FALSE
  )
  list(stats = stats, audit = audit)
}

run_fgsea <- function(result_file, label) {
  prepared <- prepare_stats(result_file, label)
  set.seed(20260523)
  res <- fgseaMultilevel(
    pathways = pathways,
    stats = prepared$stats,
    minSize = 15,
    maxSize = 500,
    eps = 0
  )
  res <- as.data.table(res)
  res[, direction := ifelse(NES > 0, "Higher in severe undernutrition", "Lower in severe undernutrition")]
  res[, leadingEdge := vapply(leadingEdge, paste, collapse = "; ", FUN.VALUE = character(1))]
  setorder(res, padj, pval)
  fwrite(res, file.path(out_dir, paste0("reactome_fgsea_", label, ".tsv")), sep = "\t")
  list(result = res, audit = prepared$audit)
}

full <- run_fgsea(full_file, "all_27_samples")
sens <- run_fgsea(sens_file, "sensitivity_excluding_102-00459-B")
mapping_audit <- rbind(full$audit, sens$audit)
fwrite(mapping_audit, file.path(out_dir, "fgsea_identifier_mapping_audit.tsv"), sep = "\t")

combined <- merge(
  full$result[, .(pathway, full_NES = NES, full_FDR = padj, full_direction = direction)],
  sens$result[, .(pathway, sensitivity_NES = NES, sensitivity_FDR = padj, sensitivity_direction = direction)],
  by = "pathway"
)
combined[, same_direction := sign(full_NES) == sign(sensitivity_NES)]
combined[, stable_FDR_lt_0.05 := same_direction & full_FDR < 0.05 & sensitivity_FDR < 0.05]
combined[, stable_FDR_lt_0.10 := same_direction & full_FDR < 0.10 & sensitivity_FDR < 0.10]
setorder(combined, -stable_FDR_lt_0.05, sensitivity_FDR, full_FDR)
fwrite(combined, file.path(out_dir, "reactome_fgsea_pathway_robustness.tsv"), sep = "\t")

summary <- data.frame(
  metric = c(
    "Full model pathways at FDR <0.05",
    "Sensitivity model pathways at FDR <0.05",
    "Stable concordant pathways at FDR <0.05",
    "Stable concordant pathways at FDR <0.10",
    "Pathway NES Spearman correlation",
    "Pathway direction agreement"
  ),
  value = c(
    sum(full$result$padj < 0.05, na.rm = TRUE),
    sum(sens$result$padj < 0.05, na.rm = TRUE),
    sum(combined$stable_FDR_lt_0.05, na.rm = TRUE),
    sum(combined$stable_FDR_lt_0.10, na.rm = TRUE),
    cor(combined$full_NES, combined$sensitivity_NES, method = "spearman", use = "complete.obs"),
    mean(combined$same_direction, na.rm = TRUE)
  )
)
fwrite(summary, file.path(out_dir, "reactome_fgsea_summary.tsv"), sep = "\t")
writeLines(capture.output(sessionInfo()), file.path(out_dir, "sessionInfo_fgsea.txt"))

cat("\nExact fgsea confirmation complete.\n")
print(summary)
cat("\nResults directory:", out_dir, "\n")
```
