## Supplementary Data S1 for "Distinct transcriptional remodeling of immune and metabolic pathways in severe undernutrition during latent tuberculosis infection does not consistently predict tuberculosis progression across independent cohorts": 03_refresh_activeTB_gate_exact_fgsea.R

```
#!/usr/bin/env Rscript
# Refresh the internal active-TB convergence gate against the exact-confirmed
# GSE152218 undernutrition programme generated by canonical limma-voom/fgsea.

suppressPackageStartupMessages({
  library(edgeR)
  library(limma)
  library(fgsea)
  library(data.table)
})

args <- commandArgs(trailingOnly = TRUE)
root <- if (length(args) >= 1) normalizePath(args[1], mustWork = FALSE) else normalizePath(".", mustWork = FALSE)
source(file.path(root, "scripts/R/phaseB_helpers.R"))

count_file <- file.path(root, "data/raw/GSE152218_features_combined.txt.gz")
meta_file <- file.path(root, "data/metadata/gse152218_severe_undernutrition_TB_axis_metadata.tsv")
map_file <- file.path(root, "data/mapping/gencode.v49.ENSG_to_symbol.tsv.gz")
gmt_file <- file.path(root, "data/pathways/ReactomePathways.gmt")
gmt_zip <- file.path(root, "data/pathways/ReactomePathways.gmt.zip")
under_limma <- file.path(root, "results/R_confirmation/limma_voom/limma_voom_sensitivity_excluding_102-00459-B.tsv")
under_fgsea <- file.path(root, "results/R_confirmation/reactome_fgsea/reactome_fgsea_sensitivity_excluding_102-00459-B.tsv")
under_robust <- file.path(root, "results/R_confirmation/reactome_fgsea/reactome_fgsea_pathway_robustness.tsv")
out_dir <- file.path(root, "results/R_confirmation_phaseB/activeTB_gate")
dir.create(out_dir, recursive = TRUE, showWarnings = FALSE)

if (!file.exists(gmt_file)) {
  if (!file.exists(gmt_zip)) stop("Missing ReactomePathways.gmt(.zip).")
  unzip(gmt_zip, exdir = file.path(root, "data/pathways"))
}
required <- c(count_file, meta_file, map_file, gmt_file, under_limma, under_fgsea, under_robust)
missing <- required[!file.exists(required)]
if (length(missing)) stop("Missing required files:\n", paste(missing, collapse="\n"))

meta <- fread(meta_file, data.table = FALSE)
raw <- fread(cmd = paste("gzip -dc", shQuote(count_file)), data.table = FALSE, check.names = FALSE)
gene_id <- sub("\\..*$", "", as.character(raw[[1]]))
if (!all(meta$subject_title %in% colnames(raw))) stop("Active-TB metadata sample identifiers do not match counts.")

counts <- as.matrix(raw[, meta$subject_title, drop = FALSE])
mode(counts) <- "numeric"
rownames(counts) <- gene_id
colnames(counts) <- meta$sample_accession
rownames(meta) <- meta$sample_accession
meta <- meta[colnames(counts), , drop = FALSE]
meta$disease <- factor(meta$tb_status, levels = c("LTBI", "activeTB"))
meta$sex <- factor(meta$sex)
meta$age_z <- as.numeric(scale(as.numeric(meta$age_years)))

run_axis <- function(keep_samples, suffix) {
  mm <- meta[keep_samples, , drop = FALSE]
  y <- DGEList(counts = counts[, rownames(mm), drop = FALSE])
  keep <- rowSums(y$counts >= 10) >= 3
  y <- y[keep, , keep.lib.sizes = FALSE]
  y <- calcNormFactors(y, method = "TMM")
  design <- model.matrix(~ disease + age_z + sex, data = mm)
  pdf(file.path(out_dir, paste0("voom_mean_variance_activeTB_", suffix, ".pdf")), width=7, height=5.5)
  v <- voom(y, design, plot = TRUE)
  dev.off()
  fit <- eBayes(lmFit(v, design))
  tab <- topTable(fit, coef = "diseaseactiveTB", number = Inf, sort.by = "P")
  tab$ensembl_gene_id <- rownames(tab)
  tab$direction <- ifelse(tab$logFC > 0, "Higher in active TB", "Lower in active TB")
  tab <- tab[, c("ensembl_gene_id","logFC","AveExpr","t","P.Value","adj.P.Val","B","direction")]
  fwrite(tab, file.path(out_dir, paste0("limma_voom_activeTB_", suffix, ".tsv")), sep="\t")
  data.frame(model=suffix, samples=ncol(y), genes_tested=nrow(y),
             FDR_lt_0.05=sum(tab$adj.P.Val < 0.05, na.rm=TRUE),
             FDR_lt_0.10=sum(tab$adj.P.Val < 0.10, na.rm=TRUE))
}

summ <- rbind(
  run_axis(rep(TRUE, nrow(meta)), "all_30_samples"),
  run_axis(meta$subject_title != "102-00459-B", "sensitivity_excluding_102-00459-B")
)
fwrite(summ, file.path(out_dir, "activeTB_limma_model_summary.tsv"), sep="\t")

# Canonical active-TB Reactome fgsea from sensitivity ranking.
mapping <- fread(cmd = paste("gzip -dc", shQuote(map_file)))
mapping[, ensembl_gene_id := sub("\\..*$", "", ensembl_gene_id)]
mapping <- unique(mapping[, .(ensembl_gene_id, gene_symbol)])
axis <- fread(file.path(out_dir, "limma_voom_activeTB_sensitivity_excluding_102-00459-B.tsv"))
axis[, ensembl_gene_id := sub("\\..*$", "", ensembl_gene_id)]
axis_mapped <- merge(axis, mapping, by="ensembl_gene_id", all=FALSE)
axis_mapped <- axis_mapped[!is.na(gene_symbol) & gene_symbol != "" & !is.na(t)]
axis_mapped[, abs_t := abs(t)]
setorder(axis_mapped, -abs_t)
axis_mapped <- axis_mapped[!duplicated(gene_symbol)]
axis_stats <- axis_mapped$t
names(axis_stats) <- axis_mapped$gene_symbol
axis_stats <- sort(axis_stats, decreasing=TRUE)

pathways <- gmtPathways(gmt_file)
set.seed(20260524)
axis_fg <- as.data.table(fgseaMultilevel(pathways, axis_stats, minSize=15, maxSize=500, eps=0))
axis_fg[, direction := ifelse(NES > 0, "Higher in active TB", "Lower in active TB")]
axis_fg[, leadingEdge := vapply(leadingEdge, paste, collapse="; ", FUN.VALUE=character(1))]
setorder(axis_fg, padj, pval)
fwrite(axis_fg, file.path(out_dir, "reactome_fgsea_activeTB_sensitivity.tsv"), sep="\t")

# Gene-level exact ranking convergence.
under_gene <- fread(under_limma)
under_gene[, ensembl_gene_id := sub("\\..*$", "", ensembl_gene_id)]
gene_cmp <- merge(
  under_gene[, .(ensembl_gene_id, under_logFC=logFC, under_t=t)],
  axis[, .(ensembl_gene_id, activeTB_logFC=logFC, activeTB_t=t)],
  by="ensembl_gene_id"
)
gene_summary <- data.frame(
  metric = c("Shared genes", "Spearman correlation of logFC", "Spearman correlation of moderated t",
             "Directional agreement"),
  value = c(nrow(gene_cmp),
            cor(gene_cmp$under_logFC, gene_cmp$activeTB_logFC, method="spearman"),
            cor(gene_cmp$under_t, gene_cmp$activeTB_t, method="spearman"),
            mean(sign(gene_cmp$under_logFC) == sign(gene_cmp$activeTB_logFC)))
)
fwrite(gene_cmp, file.path(out_dir, "exact_gene_convergence_table.tsv"), sep="\t")
fwrite(gene_summary, file.path(out_dir, "exact_gene_convergence_summary.tsv"), sep="\t")

# Pathway convergence against canonical undernutrition fgsea sensitivity ranking.
under_path <- fread(under_fgsea)
under_stable <- fread(under_robust)[stable_FDR_lt_0.05 == TRUE]
path_cmp <- merge(
  under_path[, .(pathway, under_NES=NES, under_FDR=padj, under_direction=direction)],
  axis_fg[, .(pathway, activeTB_NES=NES, activeTB_FDR=padj, activeTB_direction=direction)],
  by="pathway"
)
path_cmp[, same_direction := sign(under_NES) == sign(activeTB_NES)]
path_cmp[, exact_stable_undernutrition_pathway := pathway %in% under_stable$pathway]
stable_cmp <- path_cmp[exact_stable_undernutrition_pathway == TRUE]
path_summary <- data.frame(
  metric = c("Common Reactome pathways", "Pathway NES Spearman correlation", "Pathway direction agreement",
             "Exact stable undernutrition pathways evaluated", "Exact stable pathways same direction",
             "Exact stable pathways opposite direction", "Opposite-direction stable pathways with activeTB FDR <0.05"),
  value = c(nrow(path_cmp),
            cor(path_cmp$under_NES, path_cmp$activeTB_NES, method="spearman"),
            mean(path_cmp$same_direction),
            nrow(stable_cmp),
            sum(stable_cmp$same_direction),
            sum(!stable_cmp$same_direction),
            sum(!stable_cmp$same_direction & stable_cmp$activeTB_FDR < 0.05, na.rm=TRUE))
)
setorder(path_cmp, -exact_stable_undernutrition_pathway, activeTB_FDR)
fwrite(path_cmp, file.path(out_dir, "exact_pathway_convergence_table.tsv"), sep="\t")
fwrite(stable_cmp[same_direction == FALSE][order(activeTB_FDR)],
       file.path(out_dir, "exact_stable_pathways_opposite_to_activeTB.tsv"), sep="\t")
fwrite(path_summary, file.path(out_dir, "exact_pathway_convergence_summary.tsv"), sep="\t")
write_session(out_dir, "activeTB_gate")

cat("\nExact active-TB gate refresh completed.\n")
print(summ)
cat("\nGene-level convergence summary:\n"); print(gene_summary)
cat("\nPathway-level convergence summary:\n"); print(path_summary)
```
