## Supplementary Data S1 for "Distinct transcriptional remodeling of immune and metabolic pathways in severe undernutrition during latent tuberculosis infection does not consistently predict tuberculosis progression across independent cohorts": 04_confirm_GSE79362_exact_modules.R

```
#!/usr/bin/env Rscript
# R confirmation of frozen seven-module projection into GSE79362.
# The seven representative pathways and directions are fixed by exact GSE152218 fgsea.

suppressPackageStartupMessages(library(data.table))

args <- commandArgs(trailingOnly=TRUE)
root <- if (length(args) >= 1) normalizePath(args[1], mustWork=FALSE) else normalizePath(".", mustWork=FALSE)
source(file.path(root, "scripts/R/phaseB_helpers.R"))

expr_file <- file.path(root, "data/external_validation/GSE79362/curatedTBData/GSE79362_assay_curated.tsv.gz")
meta_file <- file.path(root, "data/external_validation/GSE79362/curatedTBData/GSE79362_colData.tsv")
module_file <- file.path(root, "data/modules/exact_confirmed_seven_module_programme.tsv")
gmt_file <- file.path(root, "data/pathways/ReactomePathways.gmt")
gmt_zip <- file.path(root, "data/pathways/ReactomePathways.gmt.zip")
out_dir <- file.path(root, "results/R_confirmation_phaseB/GSE79362")
dir.create(out_dir, recursive=TRUE, showWarnings=FALSE)

if (!file.exists(gmt_file)) unzip(gmt_zip, exdir=file.path(root, "data/pathways"))
required <- c(expr_file, meta_file, module_file, gmt_file)
missing <- required[!file.exists(required)]
if (length(missing)) stop("Missing GSE79362 required input(s):\n", paste(missing, collapse="\n"))

expr_dt <- fread(cmd=paste("gzip -dc", shQuote(expr_file)))
genes <- expr_dt[[1]]
expr <- as.matrix(expr_dt[, -1, with=FALSE])
rownames(expr) <- genes
# If duplicate symbols exist, keep row means after aggregating by symbol.
expr <- rowsum(expr, group=rownames(expr), reorder=FALSE) /
  as.numeric(table(factor(rownames(expr), levels=unique(rownames(expr)))))
meta <- fread(meta_file, data.table=FALSE)
rownames(meta) <- meta$sample_id
expr <- expr[, meta$sample_id, drop=FALSE]

meta$TimeToTB_days <- parse_numeric_time(meta$TimeToTB)
meta$baseline <- meta$MeasurementTime == "000 Day(s)"
meta$progressor <- meta$Progression == "Positive"
meta$Gender <- factor(meta$Gender)
meta$Ethnicity <- factor(meta$Ethnicity)
meta$ACS_cohort <- factor(meta$ACS_cohort)
meta$Age <- as.numeric(meta$Age)

modules <- fread(module_file)
sets <- read_gmt_simple(gmt_file)

baseline_ids <- meta$sample_id[meta$baseline]
z_all <- zscore_rows(expr, reference_columns=baseline_ids)
scores <- data.table(sample_id=meta$sample_id)
coverage <- list()
for (i in seq_len(nrow(modules))) {
  gs <- intersect(sets[[modules$representative_pathway[i]]], rownames(z_all))
  scores[[modules$theme[i]]] <- colMeans(z_all[gs, , drop=FALSE], na.rm=TRUE) * modules$orientation_multiplier[i]
  coverage[[i]] <- data.frame(theme=modules$theme[i], pathway=modules$representative_pathway[i],
                              exact_direction=modules$exact_direction[i], reactome_genes=length(sets[[modules$representative_pathway[i]]]),
                              represented_genes=length(gs),
                              coverage_percent=100*length(gs)/length(sets[[modules$representative_pathway[i]]]))
}
coverage <- rbindlist(coverage)
composite <- "Frozen undernutrition programme composite"
theme_names <- modules$theme
scores[[composite]] <- rowMeans(as.data.frame(scores[, ..theme_names]), na.rm=TRUE)
dat <- merge(meta, scores, by="sample_id")

score_names <- c(composite, modules$theme)
test_cross_sectional <- function(df, outcome_name, label, adjusted=TRUE) {
  y <- as.logical(df[[outcome_name]])
  rows <- list()
  for (score in score_names) {
    cases <- df[y, score]; controls <- df[!y, score]
    wt <- wilcox.test(cases, controls, exact=FALSE)
    auc <- as.numeric(wilcox.test(cases, controls, exact=FALSE)$statistic) / (sum(y)*sum(!y))
    or_sd <- p_adj <- NA_real_
    if (adjusted) {
      d <- df[complete.cases(df[, c(score, outcome_name, "Age", "Gender", "Ethnicity", "ACS_cohort")]), ]
      d$score_z <- as.numeric(scale(d[[score]]))
      fit <- glm(as.formula(paste(outcome_name, "~ score_z + Age + Gender + Ethnicity + ACS_cohort")),
                 data=d, family=binomial())
      or_sd <- exp(coef(fit)["score_z"])
      p_adj <- summary(fit)$coefficients["score_z","Pr(>|z|)"]
    }
    rows[[score]] <- data.frame(analysis=label, score=score, case_n=sum(y), control_n=sum(!y),
                                case_median=median(cases, na.rm=TRUE), control_median=median(controls, na.rm=TRUE),
                                median_difference_case_minus_control=median(cases, na.rm=TRUE)-median(controls, na.rm=TRUE),
                                mann_whitney_p=wt$p.value, auc_undernutrition_orientation=auc,
                                adjusted_OR_per_SD=or_sd, adjusted_p=p_adj)
  }
  out <- rbindlist(rows)
  path <- out$score != composite
  out$pathway_BH_FDR <- NA_real_
  out$pathway_BH_FDR[path] <- p.adjust(out$mann_whitney_p[path], method="BH")
  out
}

baseline <- dat[dat$baseline, ]
baseline$case <- baseline$progressor
primary <- test_cross_sectional(baseline, "case", "Baseline: future progression at any follow-up", TRUE)

short <- baseline[!baseline$progressor | (baseline$TimeToTB_days > 0 & baseline$TimeToTB_days <= 365), ]
short$case <- short$progressor & short$TimeToTB_days > 0 & short$TimeToTB_days <= 365
secondary <- test_cross_sectional(short, "case", "Baseline: TB progression within 12 months", FALSE)

pred <- dat[dat$progressor & !is.na(dat$TimeToTB_days) & dat$TimeToTB_days > 0, ]
pairs <- list()
for (pid in unique(pred$PatientID)) {
  d <- pred[pred$PatientID == pid, ]
  if (nrow(d) >= 2) {
    far <- d[which.max(d$TimeToTB_days), ]
    near <- d[which.min(d$TimeToTB_days), ]
    rec <- data.frame(PatientID=pid, far_days_to_TB=far$TimeToTB_days, near_days_to_TB=near$TimeToTB_days)
    for (score in score_names) {
      rec[[paste0(score, "__far")]] <- far[[score]]
      rec[[paste0(score, "__near")]] <- near[[score]]
      rec[[paste0(score, "__delta")]] <- near[[score]] - far[[score]]
    }
    pairs[[pid]] <- rec
  }
}
pairs <- rbindlist(pairs)
traj <- list()
for (score in score_names) {
  delta <- pairs[[paste0(score, "__delta")]]
  w <- wilcox.test(delta, mu=0, paired=FALSE, exact=FALSE)
  traj[[score]] <- data.frame(score=score, paired_progressors=nrow(pairs),
                              median_delta_near_minus_far=median(delta, na.rm=TRUE),
                              wilcoxon_p=w$p.value)
}
trajectory <- rbindlist(traj)
path <- trajectory$score != composite
trajectory$pathway_BH_FDR <- NA_real_
trajectory$pathway_BH_FDR[path] <- p.adjust(trajectory$wilcoxon_p[path], method="BH")

fwrite(coverage, file.path(out_dir, "exact_module_coverage_GSE79362.tsv"), sep="\t")
fwrite(primary, file.path(out_dir, "baseline_any_future_progression_tests_R.tsv"), sep="\t")
fwrite(secondary, file.path(out_dir, "baseline_12month_progression_tests_R.tsv"), sep="\t")
fwrite(trajectory, file.path(out_dir, "progressor_trajectory_tests_R.tsv"), sep="\t")
fwrite(pairs, file.path(out_dir, "progressor_paired_scores_R.tsv"), sep="\t")
write_session(out_dir, "GSE79362")

cat("\nGSE79362 exact-module projection R confirmation completed.\n")
print(primary[score == composite])
print(secondary[score == composite])
print(trajectory[score == composite])
```
