## Supplementary Data S1 for "Distinct transcriptional remodeling of immune and metabolic pathways in severe undernutrition during latent tuberculosis infection does not consistently predict tuberculosis progression across independent cohorts": 05_confirm_GC6_portability_gated_analysis.R

```
#!/usr/bin/env Rscript
# R confirmation of GC6/GSE94438 assay-portability gate and four-module sensitivity analysis.
# The gate is technical and independent of progression outcomes.

suppressPackageStartupMessages(library(data.table))

args <- commandArgs(trailingOnly=TRUE)
root <- if (length(args) >= 1) normalizePath(args[1], mustWork=FALSE) else normalizePath(".", mustWork=FALSE)
source(file.path(root, "scripts/R/phaseB_helpers.R"))

expr_gc6_file <- file.path(root, "data/external_validation/GSE94438/curatedTBData/GSE94438_assay_curated.tsv.gz")
meta_file <- file.path(root, "data/external_validation/GSE94438/curatedTBData/GSE94438_colData.tsv")
expr_793_file <- file.path(root, "data/external_validation/GSE79362/curatedTBData/GSE79362_assay_curated.tsv.gz")
module_file <- file.path(root, "data/modules/exact_confirmed_seven_module_programme.tsv")
gmt_file <- file.path(root, "data/pathways/ReactomePathways.gmt")
gmt_zip <- file.path(root, "data/pathways/ReactomePathways.gmt.zip")
out_dir <- file.path(root, "results/R_confirmation_phaseB/GC6")
dir.create(out_dir, recursive=TRUE, showWarnings=FALSE)
if (!file.exists(gmt_file)) unzip(gmt_zip, exdir=file.path(root,"data/pathways"))
required <- c(expr_gc6_file, meta_file, expr_793_file, module_file, gmt_file)
missing <- required[!file.exists(required)]
if (length(missing)) stop("Missing GC6 required input(s):\n", paste(missing, collapse="\n"))

read_expr <- function(path) {
  x <- fread(cmd=paste("gzip -dc", shQuote(path)))
  genes <- x[[1]]
  mat <- as.matrix(x[, -1, with=FALSE])
  rownames(mat) <- genes
  rowsum(mat, group=rownames(mat), reorder=FALSE) /
    as.numeric(table(factor(rownames(mat), levels=unique(rownames(mat)))))
}
gc6_expr <- read_expr(expr_gc6_file)
g793_expr <- read_expr(expr_793_file)
meta <- fread(meta_file, data.table=FALSE)
rownames(meta) <- meta$sample_id
gc6_expr <- gc6_expr[, meta$sample_id, drop=FALSE]
meta$TimeFromExposure_months <- parse_numeric_time(meta$TimeFromExposure)
meta$TimeToTB_months <- parse_numeric_time(meta$TimeToTB)
meta$progressor <- meta$Progression == "Positive"
meta$labelled <- meta$Progression %in% c("Positive","Negative")
meta$Gender <- factor(meta$Gender)
meta$GeographicalRegion <- factor(meta$GeographicalRegion)
meta$Age <- as.numeric(meta$Age)
meta$months_sample_to_TB <- meta$TimeToTB_months - meta$TimeFromExposure_months
meta$prediagnosis_case <- meta$progressor & !is.na(meta$months_sample_to_TB) & meta$months_sample_to_TB > 0

modules <- fread(module_file)
sets <- read_gmt_simple(gmt_file)
portability <- list()
for (i in seq_len(nrow(modules))) {
  gs <- sets[[modules$representative_pathway[i]]]
  in793 <- length(intersect(gs, rownames(g793_expr)))
  ingc6 <- length(intersect(gs, rownames(gc6_expr)))
  retention <- 100*ingc6/max(in793, 1)
  retain <- retention >= 60 & ingc6 >= 20
  portability[[i]] <- data.frame(theme=modules$theme[i], representative_pathway=modules$representative_pathway[i],
                                 exact_direction=modules$exact_direction[i], orientation_multiplier=modules$orientation_multiplier[i],
                                 GSE79362_represented_genes=in793, GC6_represented_genes=ingc6,
                                 retention_vs_GSE79362_percent=retention,
                                 portability_gate=ifelse(retain, "Retained", "Excluded"))
}
portability <- rbindlist(portability)
portable <- portability[portability_gate == "Retained"]
if (nrow(portable) != 4) stop("Expected four technically portable GC6 modules; observed ", nrow(portable))

ids <- meta$sample_id[meta$labelled]
z <- zscore_rows(gc6_expr[, ids, drop=FALSE], reference_columns=ids)
scores <- data.table(sample_id=ids)
for (i in seq_len(nrow(portable))) {
  gs <- intersect(sets[[portable$representative_pathway[i]]], rownames(z))
  scores[[portable$theme[i]]] <- colMeans(z[gs, , drop=FALSE], na.rm=TRUE) * portable$orientation_multiplier[i]
}
composite <- "Portable four-module undernutrition-oriented composite"
portable_theme_names <- portable$theme
scores[[composite]] <- rowMeans(as.data.frame(scores[, ..portable_theme_names]), na.rm=TRUE)
dat <- merge(meta[meta$labelled, ], scores, by="sample_id")
score_names <- c(composite, portable$theme)

test_design <- function(df, outcome_name, label) {
  y <- as.logical(df[[outcome_name]])
  rows <- list()
  for (score in score_names) {
    cases <- df[y, score]; controls <- df[!y, score]
    wt <- wilcox.test(cases, controls, exact=FALSE)
    auc <- as.numeric(wt$statistic)/(sum(y)*sum(!y))
    d <- df[complete.cases(df[, c(score, outcome_name, "Age", "Gender", "GeographicalRegion")]), ]
    d$score_z <- as.numeric(scale(d[[score]]))
    fit <- glm(as.formula(paste(outcome_name, "~ score_z + Age + Gender + GeographicalRegion")),
               data=d, family=binomial())
    rows[[score]] <- data.frame(analysis=label, score=score, case_n=sum(y), control_n=sum(!y),
                                case_median=median(cases), control_median=median(controls),
                                median_difference_case_minus_control=median(cases)-median(controls),
                                mann_whitney_p=wt$p.value, AUC_undernutrition_orientation=auc,
                                adjusted_OR_per_SD=exp(coef(fit)["score_z"]),
                                adjusted_p=summary(fit)$coefficients["score_z","Pr(>|z|)"])
  }
  result <- rbindlist(rows)
  path <- result$score != composite
  result$pathway_BH_FDR <- NA_real_
  result$pathway_BH_FDR[path] <- p.adjust(result$mann_whitney_p[path], method="BH")
  result
}

baseline <- dat[dat$TimeFromExposure_months == 0, ]
if (length(unique(baseline$PatientID)) != nrow(baseline)) stop("Month-0 GC6 design is not one sample per participant.")
baseline$case <- baseline$progressor
primary <- test_design(baseline, "case", "Month 0 baseline: any future TB progression")

base12 <- baseline[!baseline$progressor | (baseline$TimeToTB_months >= 1 & baseline$TimeToTB_months <= 12), ]
base12$case <- base12$progressor & base12$TimeToTB_months >= 1 & base12$TimeToTB_months <= 12
secondary <- test_design(base12, "case", "Month 0 baseline: progression within 12 months")

month6 <- dat[dat$TimeFromExposure_months == 6 & (!dat$progressor | dat$TimeToTB_months > 6), ]
month6$case <- month6$progressor & month6$TimeToTB_months >= 7 & month6$TimeToTB_months <= 18
m6 <- test_design(month6, "case", "Month 6 landmark: TB within next 12 months")

pred <- dat[dat$prediagnosis_case, ]
pairs <- list()
for (pid in unique(pred$PatientID)) {
  d <- pred[pred$PatientID == pid, ]
  if (length(unique(d$TimeFromExposure_months)) >= 2) {
    early <- d[which.min(d$TimeFromExposure_months), ]
    near <- d[which.min(d$months_sample_to_TB), ]
    if (early$sample_id != near$sample_id) {
      rec <- data.frame(PatientID=pid)
      for (score in score_names) {
        rec[[paste0(score, "__early")]] <- early[[score]]
        rec[[paste0(score, "__near")]] <- near[[score]]
        rec[[paste0(score, "__delta")]] <- near[[score]] - early[[score]]
      }
      pairs[[pid]] <- rec
    }
  }
}
pairs <- rbindlist(pairs)
traj <- list()
for (score in score_names) {
  delta <- pairs[[paste0(score, "__delta")]]
  w <- wilcox.test(delta, mu=0, exact=FALSE)
  traj[[score]] <- data.frame(score=score, paired_progressors=nrow(pairs),
                              median_delta_near_minus_early=median(delta), wilcoxon_p=w$p.value)
}
trajectory <- rbindlist(traj)
path <- trajectory$score != composite
trajectory$pathway_BH_FDR <- NA_real_
trajectory$pathway_BH_FDR[path] <- p.adjust(trajectory$wilcoxon_p[path], method="BH")

fwrite(portability, file.path(out_dir, "exact_module_portability_gate_R.tsv"), sep="\t")
fwrite(primary, file.path(out_dir, "month0_baseline_any_progression_R.tsv"), sep="\t")
fwrite(secondary, file.path(out_dir, "month0_baseline_12month_progression_R.tsv"), sep="\t")
fwrite(m6, file.path(out_dir, "month6_landmark_12month_progression_R.tsv"), sep="\t")
fwrite(trajectory, file.path(out_dir, "paired_prediagnostic_trajectory_R.tsv"), sep="\t")
fwrite(pairs, file.path(out_dir, "paired_prediagnostic_scores_R.tsv"), sep="\t")
write_session(out_dir, "GC6")

cat("\nGC6 portability-gated R confirmation completed.\n")
print(portability)
print(primary[score == composite])
print(secondary[score == composite])
print(m6[score == composite])
print(trajectory[score == composite])
```
