## Supplementary Data S1 for "Distinct transcriptional remodeling of immune and metabolic pathways in severe undernutrition during latent tuberculosis infection does not consistently predict tuberculosis progression across independent cohorts": phaseB_helpers.R

```
# Shared utility functions for Phase B R confirmation.
suppressPackageStartupMessages(library(data.table))

parse_numeric_time <- function(x) {
  out <- suppressWarnings(as.numeric(sub(" .*", "", as.character(x))))
  out[is.na(x) | x %in% c("", "NA", "<NA>", "---")] <- NA_real_
  out
}

bh_adjust_paths <- function(p) {
  p.adjust(p, method = "BH")
}

zscore_rows <- function(mat, reference_columns = colnames(mat)) {
  ref <- mat[, reference_columns, drop = FALSE]
  mu <- rowMeans(ref, na.rm = TRUE)
  s <- apply(ref, 1, sd, na.rm = TRUE)
  s[s == 0 | is.na(s)] <- NA_real_
  sweep(sweep(mat, 1, mu, "-"), 1, s, "/")
}

read_gmt_simple <- function(gmt_file) {
  lines <- readLines(gmt_file)
  parts <- strsplit(lines, "\t", fixed = TRUE)
  sets <- lapply(parts, function(x) x[-c(1,2)])
  names(sets) <- vapply(parts, `[[`, character(1), 1)
  sets
}

write_session <- function(out_dir, suffix) {
  writeLines(capture.output(sessionInfo()), file.path(out_dir, paste0("sessionInfo_", suffix, ".txt")))
}
```
