## Supplementary Data S1 for "Distinct transcriptional remodeling of immune and metabolic pathways in severe undernutrition during latent tuberculosis infection does not consistently predict tuberculosis progression across independent cohorts": README_Supplementary_Data_S1.txt

Supplementary Data S1 - exact R-confirmed outputs and reproducibility files

This archive supplies machine-readable outputs supporting the reported discovery,
active-TB convergence, GSE79362 projection and GC6 assay-portability-gated analyses.
It includes derived result tables, module definitions, metadata audit files, R scripts
and R session-information files. Source datasets remain available through GEO under
GSE152218, GSE79362 and GSE94438.
